# Opposing roles for lipocalins and a CD36 family scavenger receptor in apical extracellular matrix-dependent protection of narrow tube integrity

**DOI:** 10.1101/2025.07.25.666821

**Authors:** Alexandra C. Belfi, Sage G. Aviles, Rachel Forman-Rubinsky, Hasreet K. Gill, Jennifer D. Cohen, Aleksandra Nawrocka, Axelle Bourez, Pierre van Antwerpen, Patrick Laurent, Meera V. Sundaram

## Abstract

All exposed epithelial surfaces, including the walls of internal tubes, are lined by a lipid and glycoprotein-rich apical extracellular matrix (aECM) that helps shape and protect the apical domain. Secreted lipocalins are lipid transporters frequently found within apical compartments. We show that loss of the *C. elegans* lipocalin LPR-1 disrupts the assembly of another lipocalin, LPR-3, within the pre-cuticle aECM that protects and shapes the narrow excretory duct and pore tubes. LPR-1 is apically secreted and colocalizes with LPR-3 in intracellular vesicles and lysosomes, but unlike LPR-3 it does not detectably incorporate into the aECM. Forward genetic screens for *lpr-1* suppressors identified mutations in *scav-2,* which encodes a transmembrane protein of the CD36 scavenger receptor B family. Loss of *scav-2* restored LPR-3 matrix localization and suppressed the *lpr-1* tube shaping defect, as well as the tube-shaping defects of a subset of pre-cuticle mutants, but not *lpr-3* mutants. A SCAV-2 fusion accumulated at apical surfaces of interfacial epithelial tubes, including the excretory duct and pore, and both tissue-specific suppression of *lpr-1* matrix defects and tissue-specific rescue experiments support a local role for SCAV-2 within these tubes. These data demonstrate that LPR-1 and SCAV-2 have opposing effects on narrow tube integrity by altering the content and organization of that tube’s luminal aECM, possibly by acting as transporters of an LPR-3 cofactor. These results have broadly relevant implications regarding the importance of lipocalins and scavenger receptors for aECM organization and integrity of the narrowest tubes in the body.

## Introduction

An apical extracellular matrix (aECM) lines the environmentally-exposed, apical surfaces of epithelia, such as the insides of biological tubes. Examples in humans include the vascular glycocalyx, intestinal mucin lining, and lung surfactant [1–3]. Although the aECMs of different tissues vary in their specific contents, they all typically consist of a complex mix of proteoglycans, glycoproteins, and lipoproteins or lipids that together serve to shape and protect the apical surface [4,5]. Proteoglycans and mucins can form water-retaining gels that help expand tube lumens and protect against pathogen infection [6–8], while other types of aECM glycoproteins, such as Zona Pellucida (ZP) domain proteins, form fibrillar structures that shape the apical membrane [9–11]. The lipid components of aECM can also contribute to tube-shaping properties [1,12,13] while acting as a barrier against xenobiotics and desiccation [14–17]. Defects in aECM can cause tube collapse or leakage, contributing to disorders within the pulmonary or microvascular systems [12,18,19]. Despite the widespread importance of aECMs for human health, we still have a limited understanding of the roles played by most individual aECM components and how different glycoproteins and lipids are apically transported and assembled to build aECMs with different functional properties.

Lipocalins and Scavenger Receptor B proteins (SCARBs) are two well-known families of lipid transporters. Lipocalins (“fat cups”) are small, secreted, cup-shaped proteins that transport sterols, phospholipids and other hydrophobic cargoes throughout the body [20,21] and are most often found in apical/luminal compartments, such as mammalian plasma [22], urine [23,24], or tear film [25,26]. SCARBs (including mammalian CD36, SR-B1, and LIMP-2) are transmembrane proteins capable of binding and internalizing various extracellular proteins, but best known for their roles in cholesterol and fatty acid transport [27–31]. There are many reported correlations or anti-correlations between lipocalin or SCARB levels and tissue damage and disease in patient populations [29,32–34]. Potentially relevant to these observations, several recent studies in both invertebrates and mammals have suggested that lipocalins or SCARBs may affect aECM content [14,35–39]. Here we report opposing roles of a lipocalin and a SCARB in aECM-dependent tube shaping in *C. elegans*.

In the *C. elegans* embryo, external epithelia develop in the context of a pre-cuticle aECM that is eventually endocytosed and replaced by a cuticle [40,41]. The pre-cuticle is required to shape tissues during morphogenesis, and the cuticle then maintains and refines that shape. The pre-cuticle is also required to properly assemble the cuticle, and these two matrices continue to alternate across the larval molt cycles, with each new pre-cuticle and cuticle forming below the older cuticle, which is then shed at the molt (Fig. 1A). The pre-cuticle is composed of chondroitin proteoglycans and various glycoproteins, including members of the Zona Pellucida (ZP) domain and extracellular leucine-rich repeat only (eLRRon) protein families, which are also found in many mammalian matrices [11,42]. The cuticle is composed primarily of collagens. Both matrices have a modular composition across tissues, with different subsets of components found on different regions of the epidermis and in various interfacial tubes that connect with the epidermis [41,43]. One of the essential roles of pre-cuticle is to shape the developing excretory duct and pore tubes, which are very narrow (<0.5 microns wide) unicellular tubes that connect the excretory canal cell to the outside environment for fluid excretion (Fig. 1B) [44–46].

**Figure 1.**
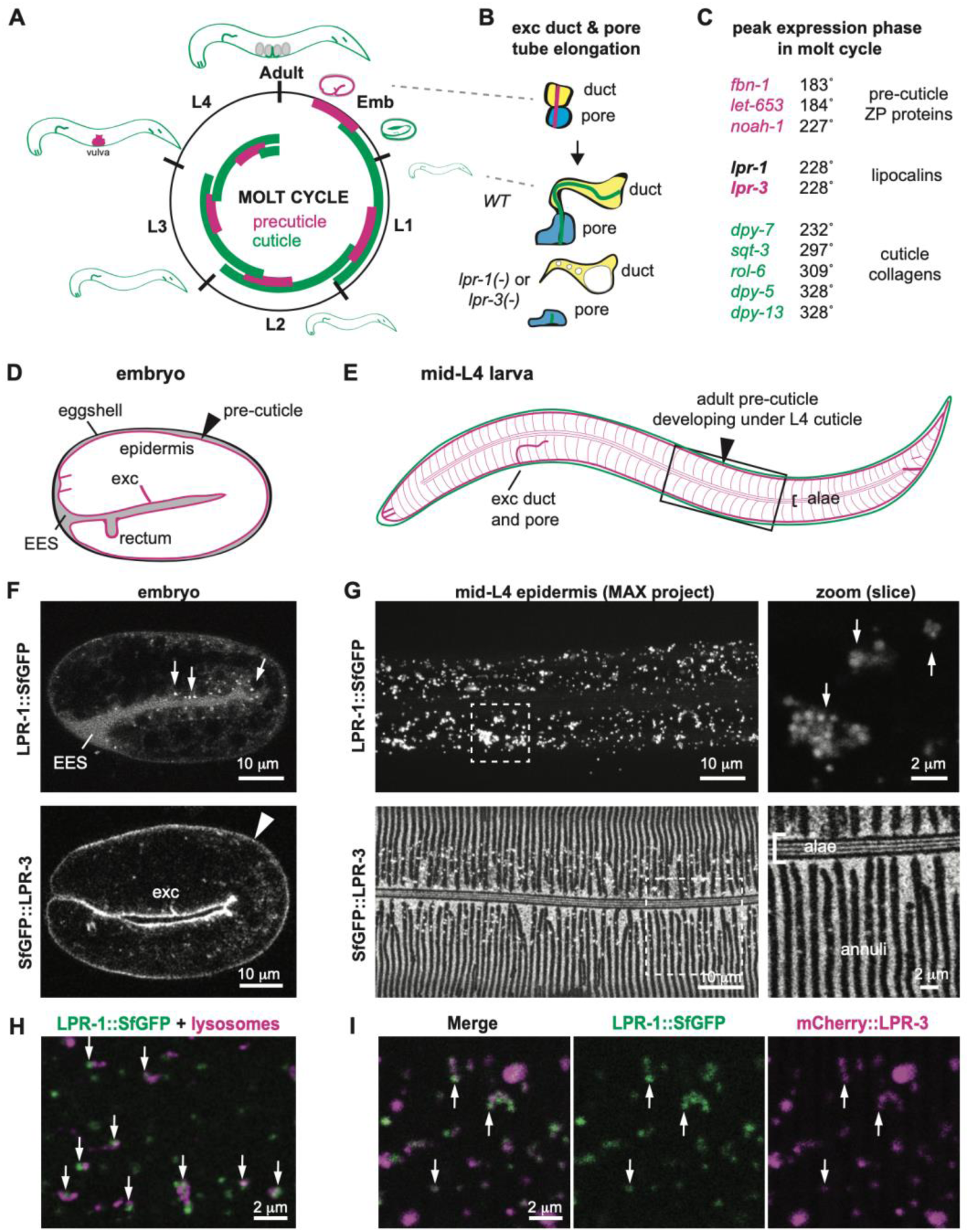
LPR-1 does not detectably incorporate into pre-cuticle but colocalizes with LPR-3 in intracellular puncta. A. The *C. elegans* molt cycle. A pre-cuticle (pink) aECM precedes each cuticle (green) and then is removed by endocytosis during cuticle assembly. Adapted from [41]m. B. Elongation of the excretory duct and pore tubes occurs during embryogenesis, when the pre-cuticle is present. In *lpr-1* or *lpr-3* mutants, much of the lumen fragments and the remaining duct lumen swells (asterisk). C. The peak oscillatory phase of *lpr-1* expression coincides with that of *lpr-3* and falls between that of known pre-cuticle components and cuticle collagens. Adapted from [51]. D. Cartoon of 2-fold embryo, a stage when pre-cuticle is present [41]. E. Cartoon of mid-L4 larva, when the adult pre-cuticle forms below the L4 cuticle. Box indicates mid-body region shown in confocal images below. F,G. Lipocalin fusion proteins in the embryo (F) and L4 larvae (G). LPR-1 fusions are found in apical puncta (arrows) and in the extraembryonic space (EES), whereas LPR-3 fusions mark the pre-cuticle matrix (arrowhead). H. Many LPR-1::SfGFP puncta were found adjacent to epidermal lysosomes or LROs marked by NUC-1::mCherry (arrows). I. LPR-1::SfGFP puncta partially overlapped with mCherry::LPR-3 puncta (arrows). Mander’s coefficient 0.72 (green+red/total green) at the L4.7 stage (n=7). All other images are representative of at least n=10 specimens imaged. See Fig. S1 for additional images.

Mutations in two *C. elegans* lipocalin-related proteins, LPR-1 and LPR-3, phenocopy pre-cuticle mutants to cause collapse of the excretory duct and pore tube lumens during embryo morphogenesis, leading to L1 larval lethality (Fig. 1B) [36,47]. Lipocalin and pre-cuticle mutants also share other matrix phenotypes such as mis-shapen cuticle ridges (alae) [36,48]. LPR-3 actually incorporates stably into the pre-cuticle, suggesting a structural role in matrix organization, but LPR-1 does not detectably bind to the matrix, and it can act tissue non-autonomously when ectopically expressed in body muscle, leaving its role unclear [47,49]. Despite these differences in protein localization, genetic double mutant analyses suggested that LPR-1 and LPR-3 function together in the same pathway [36]. Here we show that *lpr-1* loss disrupts LPR-3 matrix incorporation, and that the lethal tube defects of *lpr-1* and some pre-cuticle mutants (but not *lpr-3* mutants) can be suppressed by loss of the SCARB SCAV-2, which restores LPR-3 to the tube matrix. These results suggest that LPR-1 and SCAV-2 have opposing effects on narrow tube integrity by altering aECM contents and organization.

## Results

### LPR-1 colocalizes with LPR-3 in trafficking compartments and lysosomes

Consistent with its proposed role in pre-cuticle function, both reporter and transcriptomic data showed that *lpr-1* is most highly expressed in cuticle-producing external epithelial cells [47,50]. Furthermore, we noted that, like most pre-cuticle- and cuticle-related genes, *lpr-1* mRNA expression oscillates with the molt cycle, with its peak expression phase precisely matching that of *lpr-3* and closely following that of other pre-cuticle genes and preceding that of most cuticle collagens (Fig. 1C) [51].

To visualize endogenous LPR-1 protein during development, we used CRISPR-Cas9 to insert fluorescent tags (Superfolder (Sf) GFP or mCherry) into the endogenous *lpr-1* locus (Materials and Methods). These fusion proteins were functional based on >98% viability of the resulting animals (n>100 each). Western blotting of LPR-1::SfGFP detected a predominant band at ∼60 kd (Fig. S1), consistent with the expected full-length tagged protein. As previously observed with either N-or C-terminally tagged transgenic fusions [49], LPR-1 fusions were apically secreted prior to the 2-fold stage and accumulated in closed extracellular compartments such as the space between the embryo and the eggshell (Fig. 1D and 1F and Fig. S1). During its peak phase of expression at the mid-L4 larval stage, LPR-1::SfGFP marked numerous cytoplasmic puncta in the epidermis (Fig. 1E and 1G), most of which were found apically within the tissue (Fig. S1 and Fig. S2). LPR-1::SfGFP puncta did not overlap substantially with lipid droplets (Fig. S1) but many clustered near epidermal lysosomes or lysosome-related organelles (LROs) marked by the lysosomal deoxyribonuclease NUC-1 (Fig. 1H; Fig. S2). LPR-1 puncta disappeared in adults (Fig. S1). Most LPR-1 puncta also contained LPR-3 at the late L4 stage when LPR-3 is normally cleared from the pre-cuticle and endocytosed (Fig. 1I, Mander’s coefficient 0.72); live imaging revealed fusion of LPR-1 and LPR-3 puncta at this stage (Movie S1 and Fig. S2). Both LPR-1 and LPR-3 mCherry fusions accumulated within epidermal lysosomes marked by the SCARB SCAV-3 [52] (Fig. S2), as did control secreted mCherry fusions, consistent with the known acid-tolerance of the mCherry fluorophore [53]. These data are consistent with both LPR-1 and LPR-3 trafficking to the lysosome at the end of the molt cycle, as is typical of many pre-cuticle proteins [41].

### LPR-1 promotes LPR-3 matrix incorporation

Unlike LPR-3, LPR-1 did not detectably incorporate into the matrix at any stage examined (Fig. 1D-G and Fig. S1). Nevertheless, *lpr-1* mutant embryos had less overall accumulation of LPR-3 within the lumens of the developing excretory duct and pore tubes (Fig. 2A and B) and in the aECMs of other tissues such as the epidermis (Fig. 2E’; see below). *lpr-1* mutants had progressively increased levels of extraembryonic (non-matrix) LPR-3 (Fig. 2C-F) and more apical puncta in the epidermal cytoplasm (Fig. 2E’), indicating that secreted LPR-3 was not efficiently joining the pre-cuticle and that potentially it was being prematurely endocytosed and cleared. We conclude that LPR-1 plays an accessory role in pre-cuticle assembly, functioning upstream of LPR-3. Because *lpr-3* mutants also die due to excretory duct collapse amongst other problems [36], defects in LPR-3 matrix assembly within the excretory duct and pore can explain the lethal phenotype of most *lpr-1* mutants.

**Figure 2.**
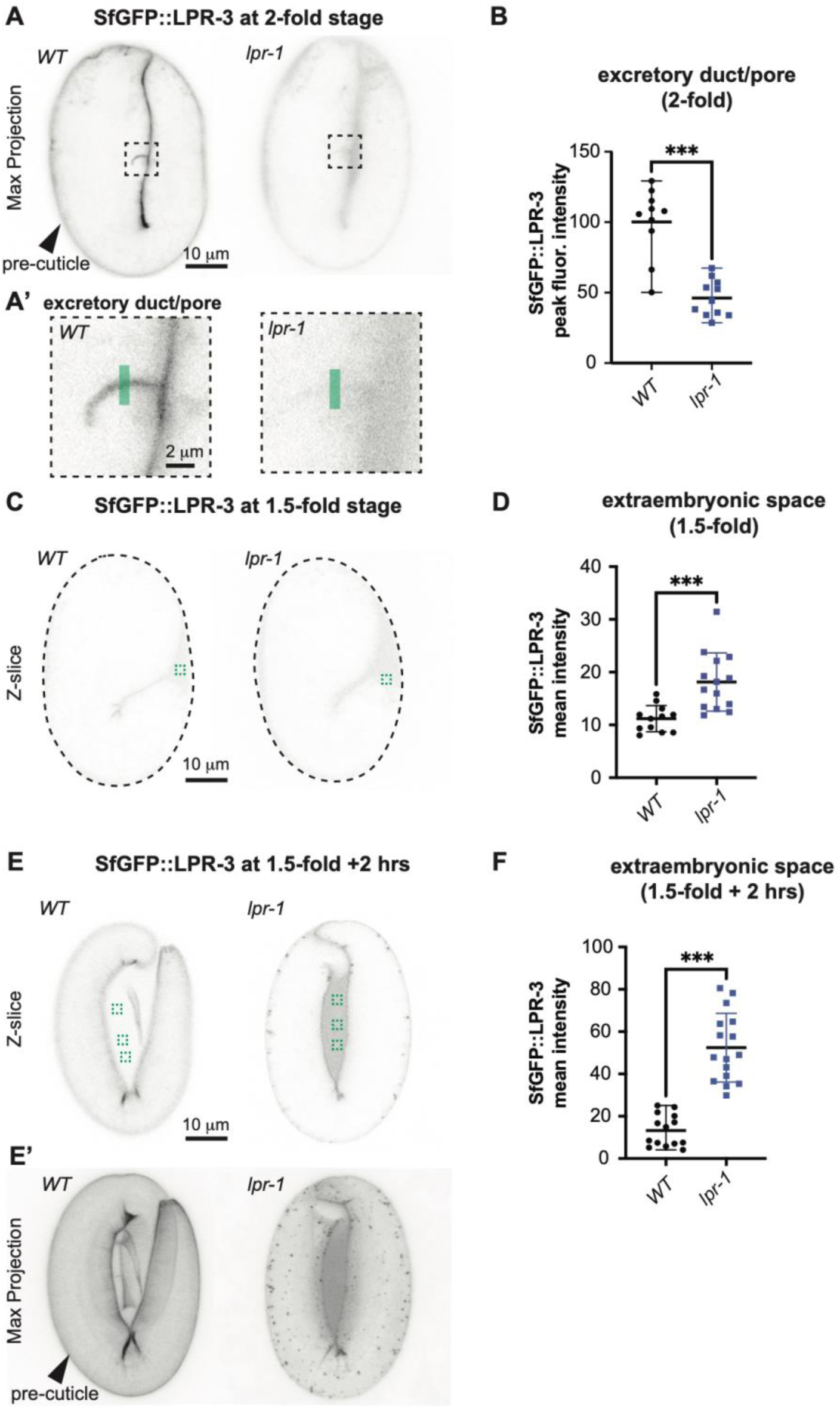
LPR-1 promotes LPR-3 localization to the pre-cuticle matrix. A,B. Loss of *lpr-1* reduces LPR-3 accumulation within the developing duct and pore tubes. A. Inverted projections of five confocal z-slices from 2-fold embryos. Boxed region is shown magnified in A’. Green line indicates duct/pore lumen region analyzed in B. B. Peak intensities of SfGFP::LPR-3 duct/pore signal as assessed using the Plot Profile tool in FIJI. ***p<0.0001, Mann-Whitney U test. C-F. *lpr-1* mutants have elevated levels of LPR-3 accumulation within the extra-embryonic space, suggesting a post-secretory defect in matrix assembly. C,E. Inverted single confocal z-slices through the medial region of 1.5-fold (C) and 1.5-fold + 2 hour (E) embryos. Dashed lines outline the embryos in C since LPR-3 signal is faint at this stage. Green box(es) indicate regions analyzed in D,F. E’. Maximum projections of the same embryos shown in E. The epidermal pre-cuticle signal also appears disorganized in *lpr-1* mutants compared to *wild-type*, and the presence of abundant puncta suggests premature LPR-3 endocytosis. D,F. Mean intensities of SfGFP::LPR-3 extra-embryonic signal as assessed using the Measure mean grey value tool in FIJI. ***p=0.0001, Mann-Whitney U test. All images are representative of at least n=10 images per genotype.

### Loss of the scavenger receptor SCAV-2 bypasses the requirement for LPR-1 and restores LPR-3 to the excretory duct/pore matrix

To better understand how LPR-1 affects the pre-cuticle matrix, we conducted an EMS mutagenesis screen to find suppressors of *lpr-1* mutant lethality. Approximately 90% of *lpr-1* mutants die as first stage (L1) larvae due to the aforementioned discontinuities in the excretory duct and pore lumens, whereas the remaining 10% maintain tube patency and survive to adulthood (Fig. 3A-C) [47,49]. After screening ∼10,000 mutagenized genomes, we identified nine suppressors that increased *lpr-1* survival to >50% (Materials and Methods, Fig. S3). Four of the best suppressors contained independent missense changes in the gene *scav-2,* and at least two of these were semi-dominant suppressors (Fig. 3D and 3E and S3). An independent *scav-2* deletion allele, *ok877*, was also a semi-dominant suppressor of *lpr-1* lethality (Fig. 3C) and restored normal duct and pore lumen morphology to *lpr-1* null mutants (Fig. 3B). These data confirm that loss of *scav-2* activity is responsible for the suppressor phenotype and demonstrate that *scav-2* is haploinsufficient and therefore dose-sensitive for that phenotype. *scav-2(ok877)* mutants did not show any obvious developmental phenotype in an *lpr-1(+)* background.

**Figure 3.**
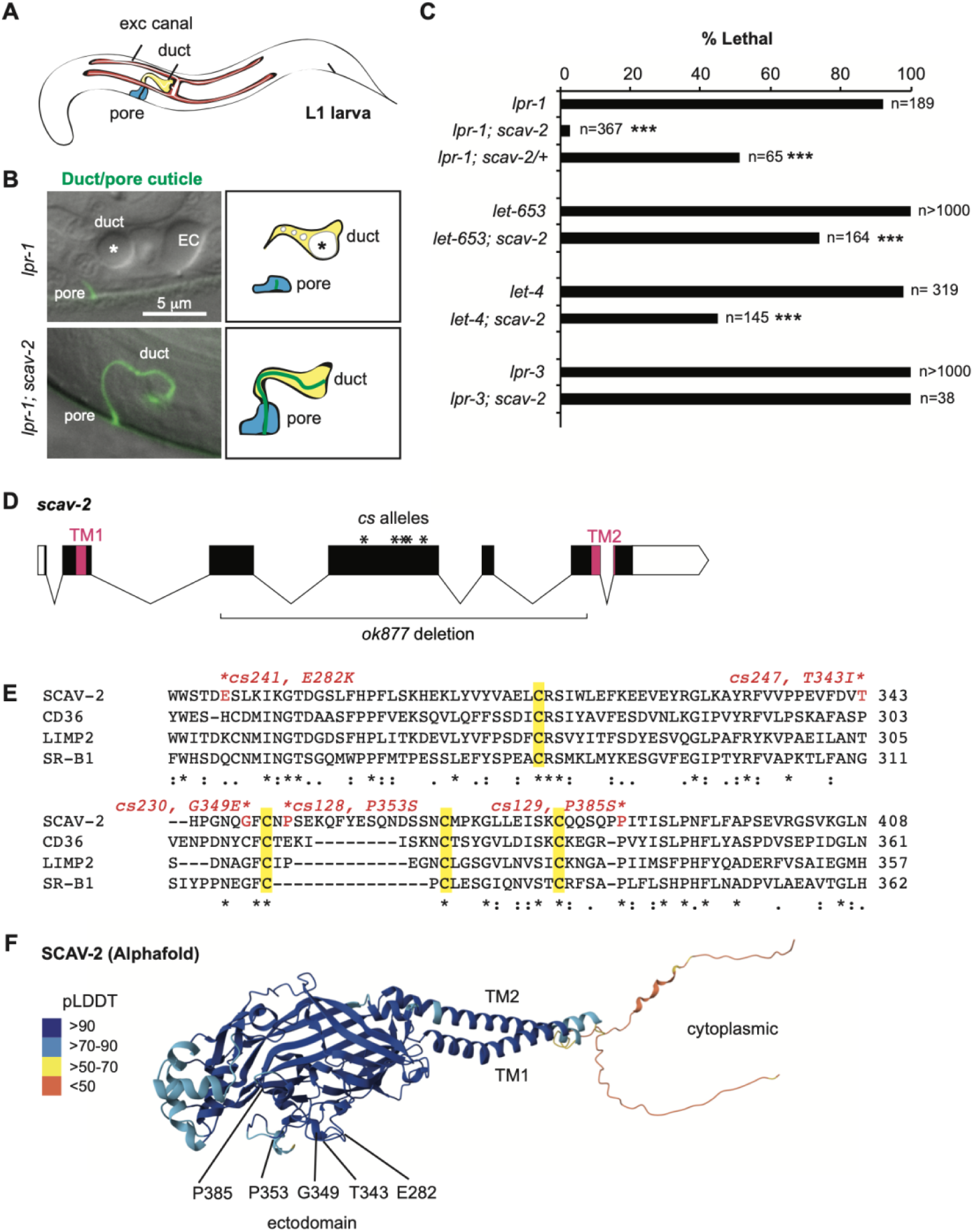
Loss of *scav-2* suppresses the lethal excretory tube defects of *lpr-1* and some pre-cuticle mutants. A. Cartoon of L1 larva, showing unicellular tubes of the excretory system. The excretory duct and pore tubes are lined by cuticle, while the upstream excretory canal cell has a different type of aECM [45]. B. Epifluorescence and Differential contrast (DIC) merged images of L1 larvae bearing an SfGFP::GRL-2 marker of the duct and pore cuticle [89], with accompanying cartoons of the cells. *lpr-1* mutants lose duct and pore tube lumen integrity [47,49] and have a large dilation but little or no cuticle in the remaining duct lumen (asterisk). Dilations are also present within the excretory canal lumen (EC). Loss of *scav-2* restored a continuous duct and pore lumen and cuticle (n=11/11), as in wild-type larvae. C. *scav-2(ok877)* partially suppressed the excretory lethal phenotype of *lpr-1(cs207), let-653(cs178)*, and *let-4(mn105)* mutants, but not *lpr-3(cs144)* mutants. ***p<0.0001, Fisher’s Exact Test, compared to *scav-2(+)* control. D. Diagram of *scav-2* locus, showing locations of molecular lesions and regions encoding both transmembrane (TM) domains. See also Fig. S3. E. Partial alignment of the SCAV-2, CD36, LIMP2, and SR-B1 ectodomains, showing conserved cysteines (yellow) and locations of *scav-2* missense lesions (red). This entire region is deleted by *ok877* (see Fig. S3). Alignment generated with Clustal Omega [90]. E. Alphafold predicted structure of SCAV-2 [91], showing positions of the *scav-2* missense alleles. The structure is colored according to the Alphafold pLDDT score, a measure of prediction confidence, which in this case is very high for most regions.

*scav-2* encodes a scavenger receptor of the SCARB family, with two predicted transmembrane domains flanking a central ectodomain, and short N- and C-terminal cytoplasmic tails (Fig. 3D and 3F). At the amino acid level, SCAV-2 is equally similar to mammalian CD36 and LIMP2 (∼28% identical) and slightly less similar to SR-B1 (∼24% identical). These three mammalian SCARBs can each bind specific lipids or lipoproteins and facilitate their transport into or across cellular membranes [29–31,54]. The central ectodomain of these receptors forms a hydrophobic tunnel structure through which lipids may access the membrane [27,55]. Interestingly, most of the *scav-2* missense alleles disrupt ectodomain residues that are identical between SCAV-2 and LIMP2 (Fig. 3E) and all cluster along one external surface of the predicted ectodomain structure (Fig. 3F), suggesting this region may be particularly important for interaction with a relevant partner or cargo. These missense alleles may be either loss of function or dominant negative; they have not been further characterized. *scav-2(ok877)* deletes most of the ectodomain and leads to a frameshift and premature stop (Fig. 3D), likely leading to nonsense-mediated decay; all further experiments were performed using this putative null allele.

While loss of *lpr-1* disrupted LPR-3 matrix localization (Fig. 2), loss of *scav-2* restored LPR-3 to the excretory/duct lumen in *lpr-1* mutants (Fig. 4A and 4C). Loss of *scav-2* did not increase LPR-3 levels in an otherwise wild-type background (Fig. 4A and 4C), suggesting that SCAV-2 does not directly transport LPR-3 but instead affects an LPR-3 partner. Nevertheless, these results indicate that LPR-1 and SCAV-2 function in opposition to respectively promote and inhibit LPR-3 matrix localization and the protection of narrow tube integrity.

**Figure 4.**
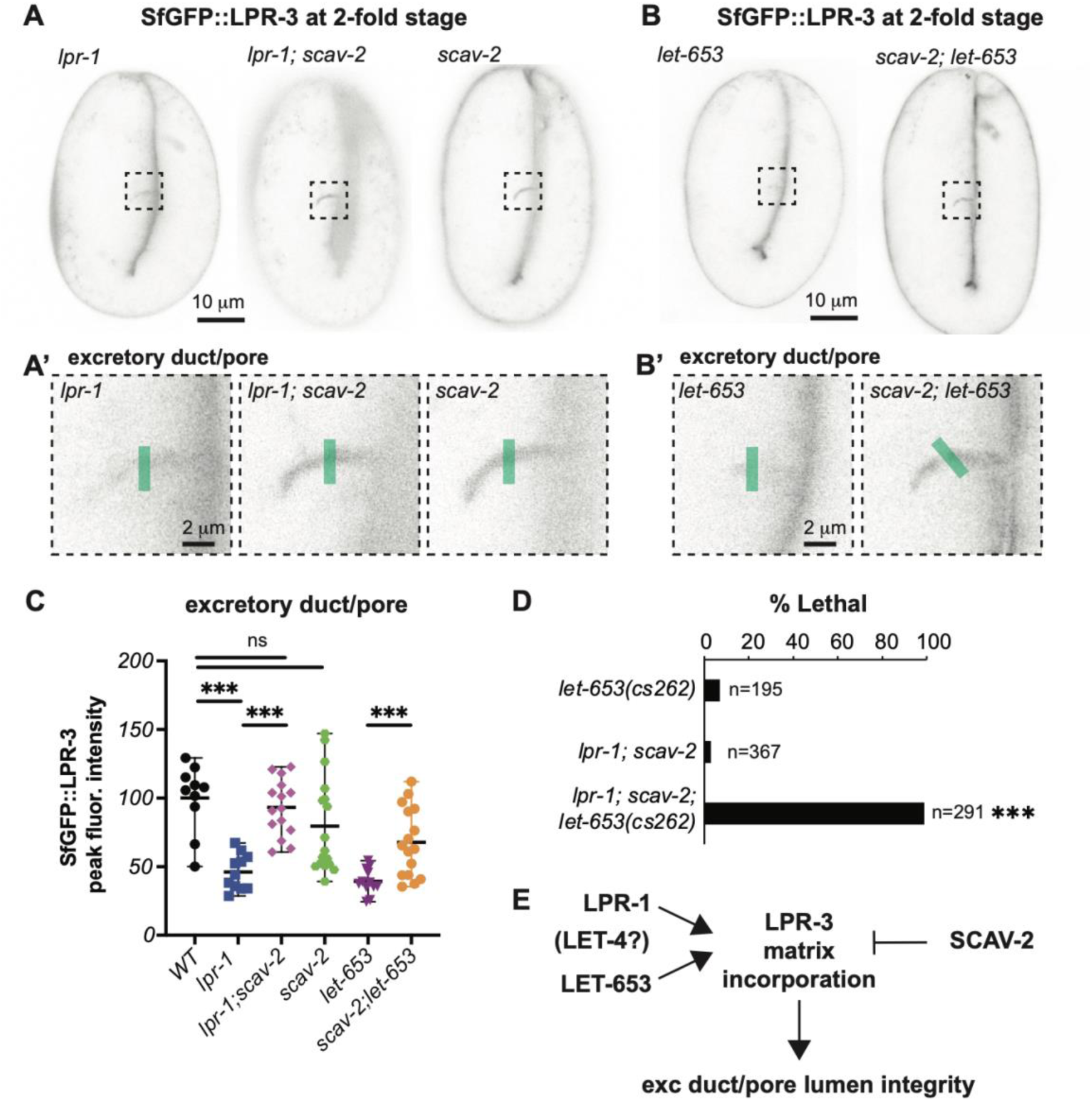
Loss of *scav-2* restores LPR-3 to the excretory duct/pore matrix of *lpr-1* and *let-653* mutants. A,B. Loss of *scav-2* restores LPR-3 accumulation within the developing duct and pore tubes. A. Inverted projections of five confocal z-slices from 2-fold embryos. Boxed region is shown magnified in A’. Green lines indicate duct/pore lumen region analyzed in C. C. Peak intensities of SfGFP::LPR-3 duct/pore signal as assessed using the Plot Profile tool in FIJI. ***p<0.0001, Mann-Whitney U test. Control data are reproduced from Fig. 2B for comparison. D. Loss of *scav-2* can’t suppress *lpr-1* lethality when *let-653* function is also partly compromised by *let-653(cs262 [LET-653::SfGFP]).* E. Genetic model of relationships.

### Loss of SCAV-2 suppresses lethality of *let-653* and *let-4* pre-cuticle mutants, but not *lpr-3* mutants

In addition to LPR-3, the ZP protein LET-653 and the eLRRon protein LET-4 are pre-cuticle components that are essential for excretory duct and pore lumen integrity [44,46]. We identified a fifth *scav-2* allele, *cs230*, in an independent EMS-mutagenesis screen for suppressors of *let-653* lethality (Fig. 3D and 3E and Fig. S3, Materials and Methods). The *scav-2*(*ok877)* null allele also partly suppressed lethality of *let-653* and *let-4* null mutants, though not of *lpr-3* mutants (Fig. 3C). Like *lpr-1* mutants, *let-653* mutants had greatly reduced LPR-3 in the excretory/duct pore lumen, but loss of *scav-2* partly restored LPR-3 to the lumen (Fig. 4B and 4C). Therefore, loss of *scav-2* ameliorates the requirements for multiple factors that act upstream of LPR-3.

Loss of LPR-1 had little or no effect on the duct/pore localization of either LET-653 or LET-4 fusions (Fig. S4), suggesting that LPR-1 acts at a step downstream or in parallel to these other matrix factors. Notably, we used a transgenic reporter for the LET-653 studies, because we could not recover any viable *lpr-1* mutants expressing an endogenously tagged LET-653::SfGFP fusion. Furthermore, most *lpr-1; scav-2*; LET-653::SfGFP larvae also died with excretory tube defects, despite the fact that most *lpr-1; scav-2* double mutants and most control LET-653::SfGFP animals are viable and healthy (Fig. 4D). This synthetic lethal interaction indicates that *lpr-1* and *let-653* act in parallel pathways and that *lpr-1* mutants are highly sensitive to even minor perturbations to LET-653 function introduced by the tag; loss of *scav-2* can compensate for loss of either the LPR-1 or LET-653 pathways, but not for simultaneous perturbations in both. We conclude that LPR-1 and LET-653 (and perhaps also LET-4) cooperate to facilitate LPR-3 assembly in the duct/pore aECM, while SCAV-2 opposes their ability to do so (Fig. 4E).

### Loss of SCAV-2 does not globally restore matrix structure to *lpr-1* mutants

In addition to affecting the excretory duct and pore matrix, *lpr-1* mutants also showed reduced and disorganized patterns of SfGFP::LPR-3 localization in epidermal pre-cuticles in both embryos and L4 larvae (Fig. 2E’ and Fig. 5A and 5B). This is consistent with prior reports that *lpr-1* mutants also have structural defects in the alae ridges of the adult cuticle [36,48]. Loss of *scav-2* did not suppress the aberrant epidermal pattern of SfGFP::LPR-3 (Fig. 5A and 5B) nor the cuticle alae defects (Fig. 5C) of *lpr-1* mutants. Therefore, loss of *scav-2* does not globally restore aECM structure, but instead specifically compensates for defects in aECM-dependent protection of the narrow excretory duct and pore tubes.

**Figure 5.**
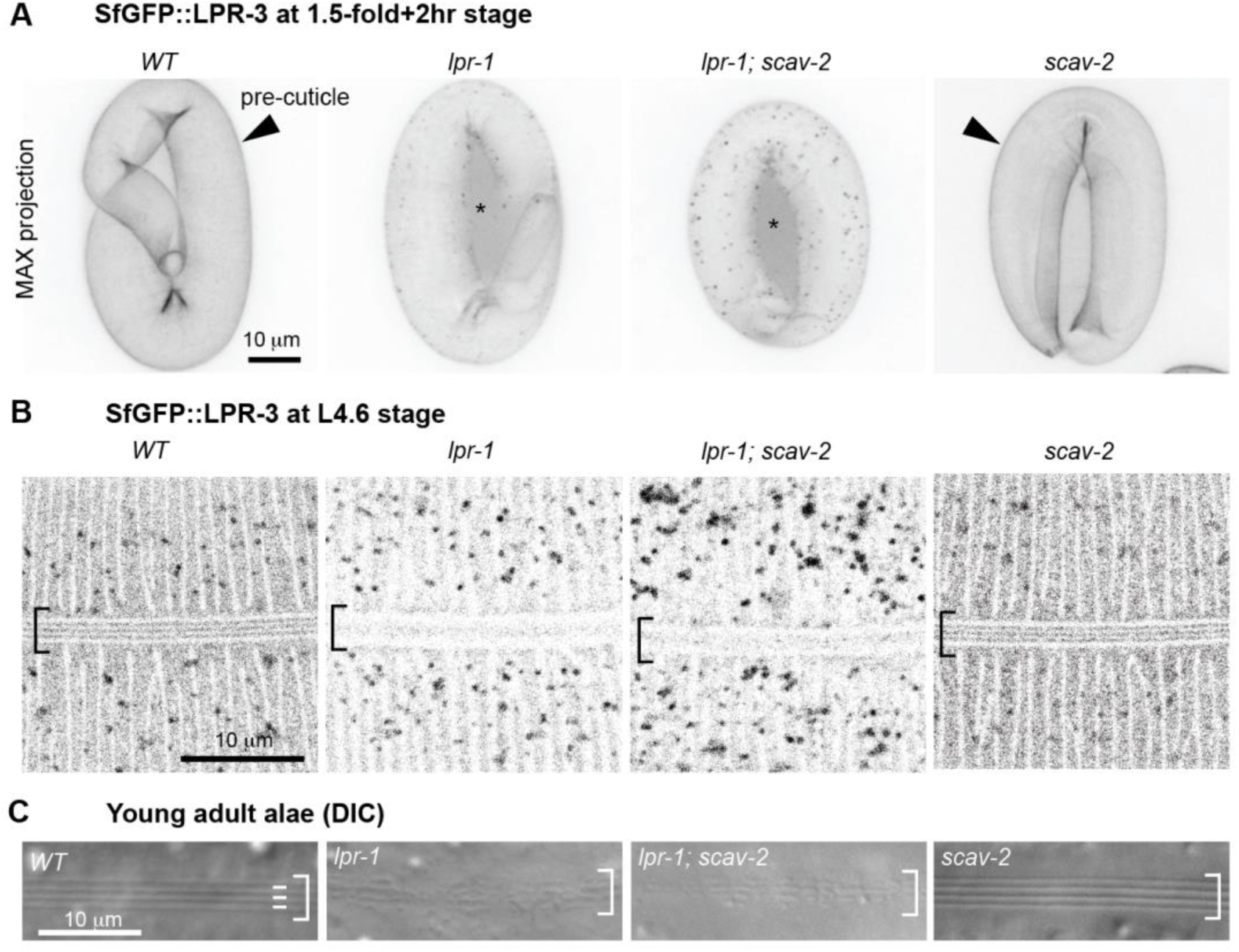
Loss of *scav-2* does not suppress *lpr-1* epidermal matrix defects. A. Inverted confocal maximum projections of SfGFP::LPR-3 in the embryonic pre-cuticle (arrowhead). *lpr-1* and *lpr-1; scav-2* mutants have reduced SfGFP::LPR-3 matrix signal and increased extra-embryonic signal (asterisk). C. Inverted confocal Z-slices of SfGFP::LPR-3 in the epidermal pre-cuticle of L4.6 stage larvae. *lpr-1* and *lpr-1; scav-2* animals have reduced SfGFP::LPR-3 matrix signal and complete loss of the organized longitudinal stripe pattern in the developing alae (brackets). C. DIC images of the alae in young adults. *scav-2* does not suppress the *lpr-1* alae defect. All images are representative of at least n=10 per genotype.

### *lpr-1* and *scav-2* mutants do not have global metabolic changes

Since lipocalins and SCARBs are best known as lipid transporters, we compared *wild-type* (*WT)*, *lpr-1* mutants, and *lpr-1; scav-2* double mutants by lipidomic profiling using liquid chromatography and mass spectrometry (LC-MS). We focused on 1.5-fold embryos for these experiments, since prior genetic rescue data indicated that LPR-1’s excretory tube-shaping function is required zygotically at or before this stage [47]. These experiments did not detect any consistent differences in lipid content among genotypes (Fig. S5). Furthermore, genetic manipulations and dietary supplementation assays failed to detect any significant effect of manipulating cholesterol or the major phospholipids phosphatidylcholine (PC) or phosphatidylethanolamine (PE) on the *lpr-1* mutant phenotype (Fig. S6). We previously reported that *lpr-1* mutants have normal epidermal barrier function [36], which depends on a lipid-rich outer cuticle layer [16,56,57]. While they don’t exclude effects on low abundance lipids or on lipid localization, as opposed to abundance (as might be predicted for lipid transporters), these experiments suggest that *lpr-1* and *scav-2* mutants do not have broad metabolic dysregulation.

### SCAV-2 acts cell autonomously in the duct and pore tubes

To ask in which cells SCAV-2 is expressed and where it is relevant, we analyzed available single cell RNA sequence (scRNAseq) data from embryos [50], generated a transcriptional reporter, and conducted tissue-specific rescue experiments. Both the scRNAseq data and the transcriptional reporter indicated that, in embryos, *scav-2* is expressed most highly in interfacial tubes, including the rectum, glial sheath and socket cells, and in the excretory duct and pore cells beginning early in tube development (Fig. 6A and 6B). Some expression was also detected in the epidermis. A *grl-2* promoter transgene driving expression of the *scav-2* cDNA only in the excretory duct and pore and several sensory glia [58] restored excretory failure and lethality to *lpr-1; scav-2* double mutants, indicating rescue of the *scav-2* suppressor phenotype (Fig. 6C). This transgene also caused more modest lethality in a wild-type background (Fig. 6C). We conclude that *scav-2* can function cell autonomously in the excretory duct and pore to oppose LPR-1 and apical ECM factors.

**Figure 6.**
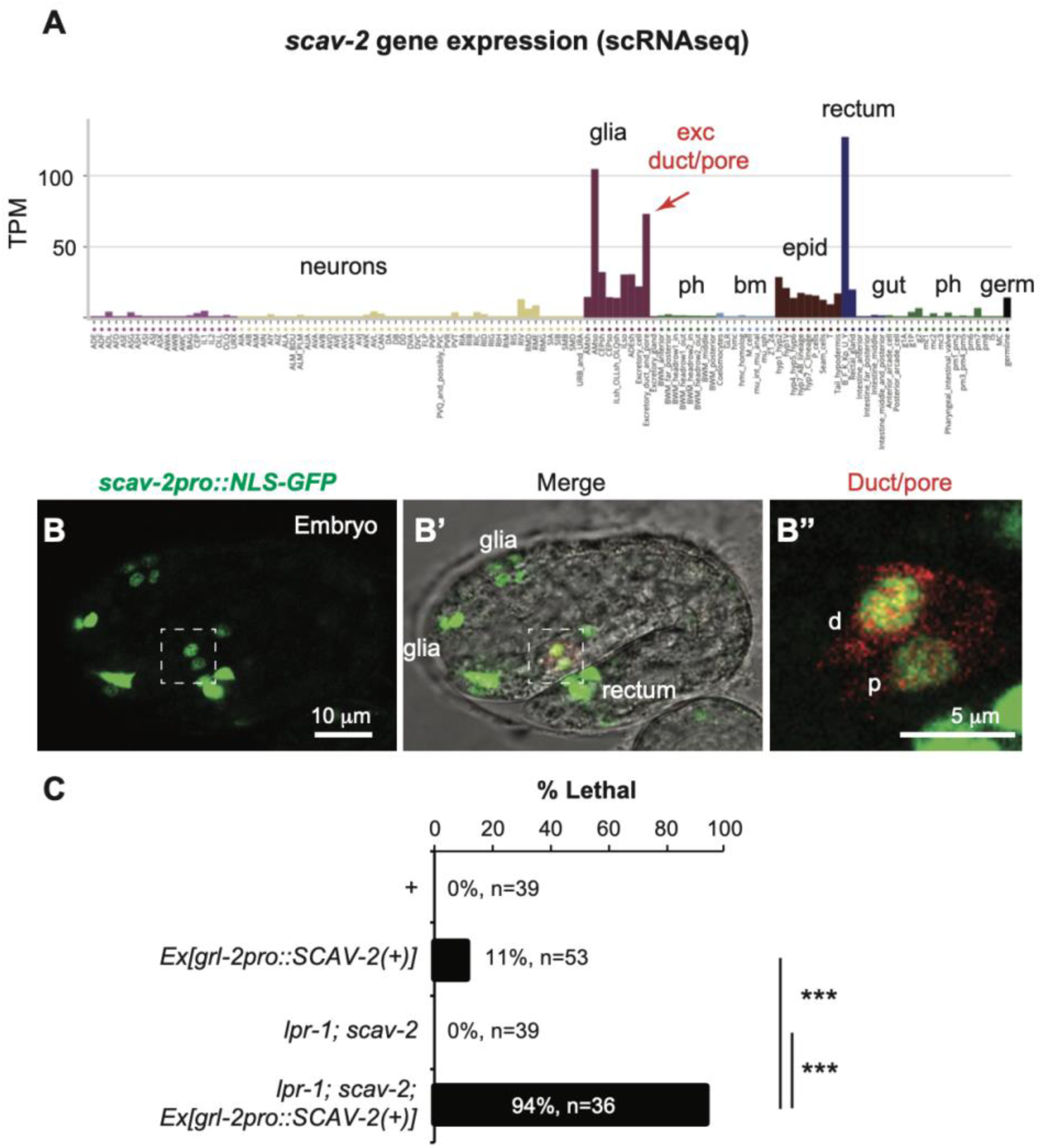
*scav-2* is expressed and functions locally within the excretory duct and pore tubes. A. The excretory duct and pore are among the cells with highest levels of *scav-2* expression. Summary of *scav-2* expression (Transcripts Per Million) in embryonic cell types, based on scRNAseq data from [50]. Plot generated with tool from ref. [92]. B. A *scav-2pro::NLS-GFP* transcriptional reporter (green) is expressed in the duct (d) and pore (p), marked by *grl-2pro::mRFP* (red) in B’,B”. Image is representative of at least n=10 embryos examined. D. A *grl-2pro::SCAV-2(+)* transgene caused minimal lethality in a wild-type background, but rescued the *scav-2* suppressor phenotype, restoring excretory defects and lethality to an *lpr-1; scav-2* strain. Transgenic and non-transgenic siblings were scored in parallel. *** n<0.001, Fisher’s Exact Test.

### SCAV-2 localizes to apical plasma membranes

SCARB proteins have been observed on both plasma membranes and internal organelle membranes, and are therefore hypothesized to transport cargo between various different compartments [27–31,37,39,52]. To assess where SCAV-2 localizes within cells *in vivo*, we used CRISPR-Cas9 to insert a fluorescent SfGFP tag at the C-terminus within the endogenous locus. The SCAV-2::SfGFP knock-in was functional based on its failure to suppress *lpr-1* defects (92% of *lpr-1; scav-2(cs256 [SCAV-2::SfGFP])* animals arrested as larvae, n=166). In both wild-type and *lpr-1* mutant backgrounds, SCAV-2::SfGFP localized to the apical membranes of the excretory duct and pore (Fig. 7A-C) and other interfacial tubes such as the rectum and vulva (Fig. 7D-G) and was observed only rarely on internal vesicles. Unlike transient pre-cuticle factors, SCAV-2 apical localization persisted throughout the molt cycle and continued into adulthood, when it became especially prominent in the vulva (Fig. 7F). We conclude that SCAV-2 is most appropriately positioned to transport materials between the apical ECM region and the cell interior.

**Figure 7:**
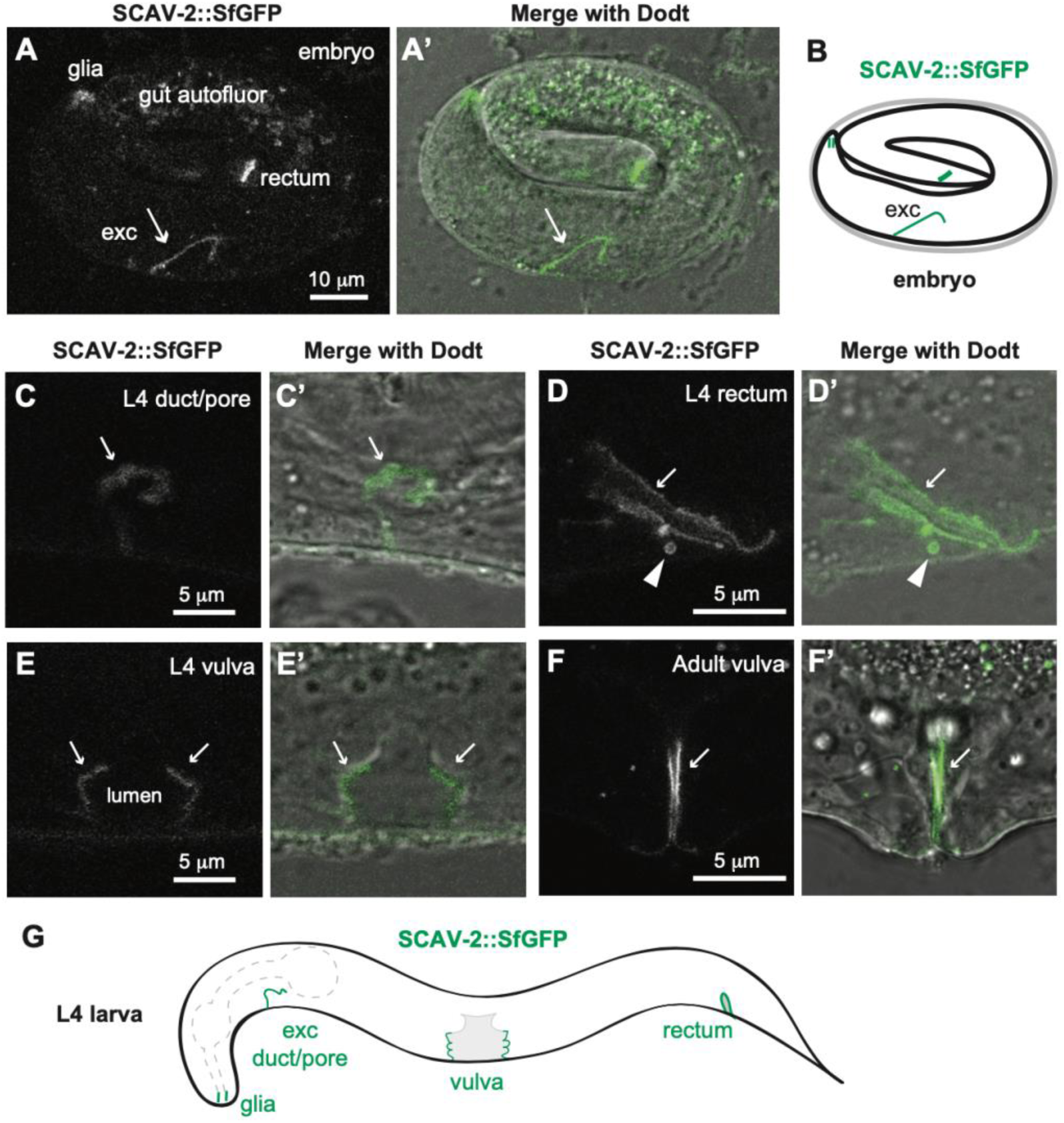
SCAV-2 localizes to apical membranes of interfacial tubes. A,A’. 3-fold embryo showing SCAV-2::SfGFP marking the apical domain of the excretory duct and pore tubes (arrow) along with the rectum and glia. B. Schematic summary of observed SCAV-2::SfGFP expression in embryos. C-E. L4 larvae showing SCAV-2::SfGFP marking the apical domains of the duct and pore (C, C’), rectum (D,D’) and vulva (E,E’) tubes. SCAV-2::SfGFP persists in the adult vulva. G. Schematic summary of observed SCAV-2::SfGFP expression at the L4 stage. Arrows indicate apical membrane signals and the arrowhead in D indicates a large intracellular vesicle. All images are representative of at least n=10 specimens imaged per stage and tissue.

## Discussion

aECMs play important roles in shaping and protecting tube lumens and are particularly critical for maintaining patency and flow in the narrowest tubes such as capillaries and alveoli [12,18,46,59–61]. This work shows that two putative lipid or lipoprotein transporters, the lipocalin LPR-1 and the CD36-related scavenger receptor SCAV-2, function in opposition to affect aECM assembly and aECM-dependent narrow tube patency in *C. elegans*. The relevant cargo(s) of these transporters are unknown but may include matrix lipids or other cofactors that contribute to LPR-3 matrix assembly. LPR-1 normally promotes the delivery of such factors to increase the LPR-3 content and tube-widening properties of aECM, while SCAV-2 normally removes such tube-widening factors from the luminal environment (Fig. 8). If mammalian lipocalins and SCARBs similarly affect aECM organization in the microvasculature and other tubular organ systems, such a role could explain many of their reported impacts on human health.

**Figure 8.**
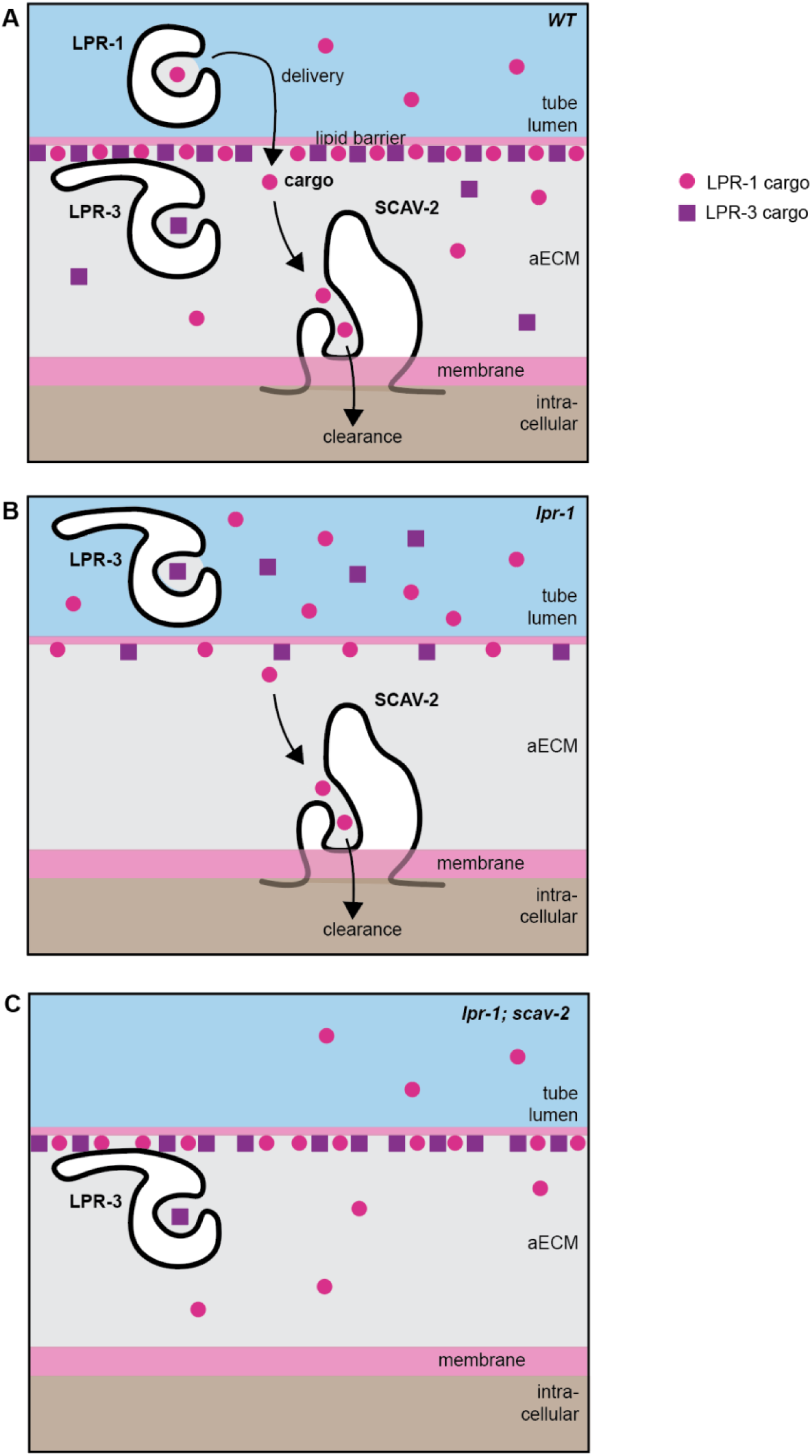
Model. A. LPR-1 and SCAV-2 function, respectively, to deliver and clear an unknown lipophilic cargo (pink circle) that helps to promote assembly or retention of LPR-3 in the aECM. It is also possible that LPR-1 and SCAV-2 transport distinct cargos that each impact LPR-3 (not shown). LPR-3 matrix association is conferred by its unique N-terminal domain [36]. LPR-3 is presumed to carry a different cargo (purple square) since the LPR-1 and LPR-3 cup domains are not interchangeable [36]. B. In the absence of LPR-1, cargo levels drop below the level needed to efficiently recruit or retain LPR-3 in the matrix. C. In the absence of both LPR-1 and SCAV-2, cargo levels are restored to homeostasis and LPR-3 can assemble in the aECM.

### Lipocalin LPR-1 promotes LPR-3 matrix incorporation

In *lpr-1* mutants, LPR-3 is still apically secreted but does not efficiently incorporate into the pre-cuticle aECM, both in the excretory duct/pore and in other parts of the body. Therefore, LPR-1 must act upstream of LPR-3 to promote its matrix delivery or incorporation (Fig. 8). LPR-1 acts in parallel to the ZP domain protein LET-653, a component of the pre-cuticle that also promotes LPR-3 matrix incorporation (Fig. 4), presumably by organizing an appropriate matrix environment.

The low abundance and non-cell autonomous action of LPR-1 are most consistent with a non-structural matrix role such as in long-distance transport or signaling [47,49]. Lipocalins can function as carriers of many different types of hydrophobic cargoes, including steroid hormones or other signaling lipids [21]. For example, mammalian ApoM transports sphingosine-1-phosphate (S1P) for signaling via the S1P receptor [62,63], while Retinol Binding Protein (RBP4) transports retinol through the bloodstream and facilitates its uptake through receptor STRA6 for eventual signaling through the retinoic acid receptor [64,65]. Although some evidence suggests that *C. elegans* LPR-1 and LPR-3 can also bind and transport S1P, mutations in the S1P pathway do not cause the lethal excretory duct phenotypes seen in *lpr-1* and *lpr-3* mutants [66]. Several essential *C. elegans* nuclear hormone receptors (NHRs) do promote matrix-relevant gene expression during the molt cycle [67,68], so LPR-1 could potentially transport a relevant NHR ligand; however, no such ligand has been identified to date and our cholesterol manipulation experiments (Fig. S6) did not support sterol involvement. An alternative model is that LPR-1 more directly delivers specific matrix lipids or lipoproteins, such as LPR-3 or an LPR-3 cofactor, that are required within the luminal environment of the excretory duct and pore tubes (Fig. 8). Studies of the other *lpr-1* suppressors (Fig. S3) may provide further clues to the LPR-1 cargo.

The frequent association of both LPR-1 and LPR-3 with epidermal lysosomes or LROs may be significant. In addition to being sites for protein degradation, lysosomes are important sites of lipid storage and metabolism [69]. Many epithelia also have specialized LROs dedicated to their particular needs; for example, lamellar bodies in the lung are specialized LROs that store, modify, and deliver lipid-rich lung surfactant [70], and *C. elegans* gut granule LROs are known sites of BODIPY-labelled lipid accumulation and ascaroside (glycolipid) biosynthesis [71–73]. *C. elegans* epidermal lysosomes or LROs are diverse in size, morphology, and spatial distribution (Fig. S2) and have been noted to change appearance over the course of the molt cycle [74]. An attractive hypothesis is that LPR-1 could be involved in transporting materials to or from a subset of these lysosomes/LROs that store and recycle aECM factors. Further studies of these compartments will be important to understand their potential functions in matrix handling.

### SCARB SCAV-2 opposes LPR-1 by promoting LPR-3 matrix removal

Scavenger receptors are so named based on their ability to remove various factors from the extracellular environment [75]. The data presented here suggest that *C. elegans* SCAV-2 acts locally to remove factors from the luminal matrix in the excretory duct and pore (Fig. 8A and 8B). Loss of *scav-2* suppresses *lpr-1* tube defects and restores LPR-3 to the luminal matrix (Fig. 8C). The SCAV-2 cargo is unlikely to be LPR-3 itself since LPR-3 matrix levels were not increased in *scav-2* single mutants; instead SCAV-2 may remove matrix lipids or other factors that function upstream to impact LPR-3 recruitment or retention (Fig. 8).

SCARB proteins are best known for their lipid and lipoprotein transport roles but could have broader functions in matrix biology. The mammalian SCARBs CD36 and SR-B1 are found on plasma membranes and are important for cellular uptake of circulating fatty acids and LDL or HDL cholesterol [76–78]. However, they can also bind other types of molecules, including aECM protein cargoes such as thrombospondin or collagen [79]. The SCARB LIMP-2 (Lysosome Integral Membrane Protein 2) localizes to endomembranes and functions both as a cholesterol transporter and as a sorting receptor that traffics specific lipid-modifying enzymes to lysosomes and LROs [30,54]; because LROs include surfactant-producing lamellar bodies, LIMP-2 can moderate the lipid content of lung surfactant [35]. A Drosophila SCARB, Emp, was recently shown to promote clearance of LDLr domain-containing proteins during maturation of the aECM within tracheal airway tubes [39]. Thus, there may be very broadly conserved roles for SCARBs in modulating aECM organization.

### Potential aECM-related roles for lipocalins and SCARBs in human disease

As mentioned in the Introduction, there are many reported correlations or anti-correlations between lipocalin or SCARB levels and tissue damage and susceptibility to infection or disease in patient populations, as well as in animal models [29,32–34]. For example, for various lipocalins, either reduced or increased levels of expression have been noted to correlate with cardiovascular disease burden and in some cases contribute to it [21], while reducing CD36 levels can either predispose or protect from cardiovascular disease depending on genetic background and diet [34]. A wide range of mechanisms have been proposed to explain these observations, most involving effects on bacterial siderophores, inflammatory signaling pathways, or lipid metabolism and storage [21,34]. Mammalian studies rarely examine the vascular aECM, which is difficult to visualize by light microscopy but whose disruption or repair could easily contribute to many of the other observed effects. The results here clearly demonstrate that both lipocalin and SCARB mutations impact aECM contents and organization in *C. elegans* and should motivate further exploration of this possibility in other systems.

## Materials and Methods

### Animal husbandry

All strains used in this study are listed in Table S1. Unless otherwise indicated, strains were grown under standard conditions on NGM plates at 20° C [80]. All experimental planning relied on data publicly available through Wormbase and the Alliance of Genome Resources [81].

### Isolation of *scav-2* alleles

Strain UP1321 [*lpr-1(cs73)*] was mutagenized with 50mM EMS [80], and P0 animals were placed on large NGM plates. After 2 weeks, plates were screened for increased population density, and individual animals were picked to establish stocks; only a single suppressed stock from each original P0 plate was kept. For each established stock, survival was quantified by collecting eggs from at least five animals and assessing survival to L4 stage three days later. We estimate that progeny from ∼5,000 F1s were screened, and a total of fourteen suppressed stocks were established, for which nine showed survival >50% and were further studied (Fig. S3). *lpr-1* suppressors were tested for linkage to chromosomes I or III by outcrossing with balancer *hT2[qIs48] (I;III)*, and then assessing what proportion of F2 non-balancer animals (*lpr-1* mutants) carried the suppressor. Four of the nine strains, which turned out to contain *scav-2* alleles, showed 100% presence of the suppressor, indicating linkage to the balancer-covered region.

Separately, strain UP2846 [*let-653(cs178) jcIs1; csIs61; csEx358 (lpr-1pro::let-653+; unc-119::gfp)*] was mutagenized with 50mM EMS and the F2 progeny screened for any surviving non-transgenic larvae. We estimate that progeny from ∼5000 F1s were screened and a single suppressed stock (with suppressor *cs230*) was identified.

The *scav-2 cs128, cs129,* and *cs230* lesions were identified by whole genome sequencing of Nextera libraries on an Illumina sequencer, followed by bioinformatic analysis with Cloudmap [82]. The *scav-2 cs241* and *cs247* lesions were identified (and the other lesions confirmed) by Sanger sequencing of PCR-amplified genomic fragments.

### *scav-2* transgenic reporters and rescue constructs

All plasmids for transgenic experiments were based on standard vector backbone pPD49.26 (Addgene), with promoter sequences cloned into MCSI and the gene of interest cloned into MCSII. See Table S2 for details.

### *lpr-1* and *scav-2* endogenous reporter fusions

Fluorescent tags were inserted into the 3’ end of the endogenous *lpr-1* locus using CRISPR-Cas9 and the self-excising cassette method [83] (see Table S2). After marker excision, edits were confirmed by Sanger sequencing.

An SfGFP fluorescent tag was inserted into the 3’ end of the endogenous *scav-2* locus using CRISPR-Cas9 and the scarless editing protocol of [84], modified by using two crRNAs simultaneously. Cas9-NLS protein was purchased from the QB3 MacroLab core at the University of California, Berkeley. Ultramers, tracr RNA and crRNAs (Table S2) were purchased from IDT. Edits in F2s were identified by visual screening for fluorescence and were verified by Sanger sequencing.

### Microscopy

Confocal images were captured with a Leica TCS SP8 or Leica TCS DMi8 confocal microscope and Leica Las X Software. All fluorescent images were captured through an HC PL APO 63x (Numerical Aperture 1.3). Z stacks were collected with 0.33 μm slices, with a 600 or 1000 Hz scanning speed and a pinhole setting of 1 AU. SfGFP fusions were excited via a 488 nm laser and emissions between 493 and 547 nm were detected using a HyD sensor. mCherry fusion proteins were excited by a 552nm laser and emissions between 583 and 784 nm were detected by a HyD sensor. Laser power (1-3%), line accumulation (1-6x) and gain settings (15-75%) varied based on fusion protein. Offset settings were not utilized. Worms were immobilized with 10 mM levamisole in M9 buffer and mounted on 2% agarose pads supplemented with 2.5% sodium azide. Embryos were staged based on minutes of development past 1.5-fold [40]. L4 larvae were staged based on vulva morphology [85,86].

### Image analysis

Images were processed using ImageJ [87]. Mander’s coefficients of overlap between LPR-1::SfGFP and mCherry::LPR-3 were obtained using the JaCoP plugin and a single confocal Z-slice from each L4.7 stage animal. Threshold settings were 51 for LPR-1::SfGFP and 43 for mCherry::LPR-3. Peak intensities of fusion proteins in the excretory duct/pore lumen were assessed using the Plot Profile tool and a 10 pixel wide line drawn across the lumen within maximum projections of five confocal Z-slices per embryo. Mean gray values of extra-embryonic fusion proteins were assessed using the Measure tool and 2 μM^2^ boxes drawn within a single confocal Z-slice per embryo. LPR-1::SfGFP puncta number in L4 animals was assessed using the Analyze Puncta tool and a threshold setting of 40. Graphs were generated using Prism/GraphPad®, where each individual dot represents a single animal. All quantitative image data were compared between genotypes using a non-parametric Mann-Whitney U-test.

### Western blots

Western blots with mixed stage LPR-1::SfGFP animals (strain UP3665) were performed as in [88] using primary antibody goat anti-GFP (Rockland Immunochemicals, 600-101-215, 1:1000 dilution) and secondary antibody anti-goat-HRP (Rockland Immunochemicals, 605–4302, 1:10,000 dilution).

### Lipidomics using liquid chromatography and mass spectrometry

Lipidomics was performed on 1.5-fold-stage embryos from strains N2 (wild type), UP2486 (*lpr-1(cs207)*), and UP3167 (*lpr-1(cs207); scav-2(ok877)*), grown in parallel on 9 cm NGM plates seeded with a thick lawn of OP50. For each strain, 3–5 biological replicates were collected across three independent experiments conducted on different days. Eggs were purified from well-fed Day1/Day2 adults as described in [16]. Briefly, eggs prepared by bleaching the adults were washed three times with M9 buffer before being resuspended in 6 mL M9 and gently shaken for 5 hours at 20 °C to allow development to the 1.5-fold stage. Embryos were purified in a 30% sucrose gradient, washed in Milli-Q water, and the pellet was snap-frozen in 2 mL tubes in liquid nitrogen. Samples were stored at –80 °C until further processing. For homogenization, frozen samples were thawed on ice, and a mixture of 0.7 mm zirconia beads (BioSpec Products, #11079107ZX) and polar solvent (500 µL Milli-Q water and 500 µL methanol) was added. Samples were then homogenized using a Bullet Blender for 5 minutes at 30 Hz. Subsequently, chloroform was added to reach a final volume of 1 mL. Samples were thoroughly mixed and centrifuged for 10 minutes at 20,000 × g to induce phase separation. After centrifugation, the apolar (chloroform) phase was collected for lipidomic analysis. This apolar phase was then dried under nitrogen gas and reconstituted in 100 µL chloroform/methanol (2:1, v/v) before injection. Reverse-phase liquid chromatography was performed using a Zorbax XDB C18 column (50 × 4.6 mm, 1.8 µm) maintained at 40 °C. The mobile phases consisted of 5 mM ammonium acetate buffer at pH 5 (solvent A) and isopropanol (solvent B). A gradient elution was applied as follows: 0 min, 90:10 (A:B); 10 min, 20:80; 25 min, 20:80; 27 min, 90:10; 30 min, 90:10, at a flow rate of 0.5 mL/min. Mass spectrometry was carried out on an Agilent QTOF 6520 instrument in autoMSMS mode with a fixed collision energy of 25 eV. Source parameters were: drying gas temperature 350 °C, gas flow 7 L/min, nebulizer pressure 50 psi, capillary voltage 4500 V, and fragmentor voltage 210 V. MS and MS/MS spectra were recorded over an m/z range of 100–1700. The acquired MS data underwent a rigorous analysis, encompassing Principal Component Analysis (PCA), Partial Least Squares Discriminant Analysis (PLS-DA), heatmaps, and univariate analysis. None of these analyses reported consistent differences between genotypes.

Subsequently, the obtained Agilent format “.d” data were converted to .mzXML format using ProteoWizard MSConvert software (version 3.03.9393, 64-bit) with the Peak Picking filter option. Preprocessing, which included peak detection, identification, grouping and smoothing, retention time correction, filtration, integration, and signal drift correction, was conducted using the freely available Galaxy Workflow4Metabolomics (W4M) platform.

### Dietary lipid supplementation experiments

5% stock solutions of egg phosphatidylcholine (Sigma: P3556), lyso-phosphatidylcholine (Sigma: L4129), or phosphatidylethanolamine (Sigma: P7943), were made with 100% ethanol and stored at −20°C. Lipid stocks were freshly diluted to 0.01% in M9 buffer just before use. For each lipid, 200 ul were applied to NGM plates and allowed to dry for several hours before seeding with normal OP50 bacteria. For cholesterol experiments, NGM plates were prepared without cholesterol or with 5x the normal amount of cholesterol. For *lpr-1* survival tests, adult P0 hermaphrodites were placed on lipid plates, F1 survivors were used for synchronized egg-lays (still on lipid plates), and F2 progeny were scored for viability. For *pcyt-1* fertility tests, adult P0 hermaphrodites were placed on lipid plates, F1 progeny were transferred as L4s to 25° C, and F2 progeny were counted to assess fertility.

## Acknowledgements

We gratefully acknowledge Juan Santana and Kevin Cullison, who performed the original screen for *lpr-1* suppressors that identified *scav-2* alleles, and Pu Pu, Lydia Malackiel, and Danielle Tsougarakis, who assisted with some experiments. We also thank Nathalie Pujol and members of our laboratory for helpful discussions and comments on the manuscript, and Andrea Stout and the CDB microscopy core for training and assistance with confocal microscopy. Some strains were obtained from the *Caenorhabditis* Genetics Center (CGC), which is funded by NIH Office of Research Infrastructure Programs P40 OD10440. This work was funded by grants from the National Institutes of Health (NIH R01GM125959 and R35GM136315 to M.V.S.) and the Belgian Fonds de la Recherche Scientifique (FNRS PDR 40020876 to P.L. and FRIA 1.E.098.24F to A.N.).

## Notes

### Competing Interest Statement

The authors have declared no competing interest.

