## Supplemental materials for "Opposing roles for lipocalins and a CD36 family scavenger receptor in apical extracellular matrix-dependent protection of narrow tube integrity"

**Supplementary Data**

Table S1: worm strain list

Table S2: plasmid and primer lists

Figures S1-S6

Movie S1

**Table S1: Strains used**

| Strain | Genotype | Source |
|---|---|---|
| N2 | *wild type* | |
| ML2578 | *let-4(mc95 [LET-4::GFP]) X* | Vuong-Brender et al 2017 |
| PHX7064 | *grl-2(syb7064 [SfGFP::GRL-2]) V* | Serra et al 2024 |
| RB973 | *scav-2(ok877) III* | CGC |
| RB1737 | *pld-1(ok2222) II* | CGC |
| UP1787 | *let-4(mn105)x; csEx173(C44H4, sur-5::gfp)* | Mancuso et al 2012 |
| UP2436 | *lpr-1(cs207) I* | Pu et al 2017 |
| UP3167 | *lpr-1(cs207) I; scav-2(ok877) III* | This work |
| UP3205 | *csEx683 [grl-2pro::scav-2(cDNA), lin-48::mRFP, RDY-2::GFP]* | This work |
| UP3210 | *lpr-1(cs207) I; scav-2(ok877) III; csEx683 [grl-2pro::scav-2(cDNA), lin-48::mRFP, RDY-2::GFP]* | This work |
| UP3244 | *let-653(cs178); csEx358[lpr-1p::LET-653b, unc-119p::GFP]* | Gill et al 2016 |
| UP3261 | *scav-2(ok877)III; lpr-3(cs144) X; csEx436[lpr-3 fosmid+myo-2::mCherry]* | This work |
| UP3330 | *csEx757[scav-2pro(5.8k)::NLS::GFP;grl-2pro::mRFP]* | This work |
| UP3422 | *csIs66 [let-653pro::SfGFP::LET-653(ZP), let-653pro::PH-mCherry, lin-48pro::mRFP] X* | Gill et al 2016 |
| UP3435 | *+/hT2[qIs48] (I;III)* | This work |
| UP3440 | *scav-2(ok877) III; let-4(mn105) IV* | This work |
| UP3561 | *pcyt-1(et9) X* | This work |
| UP3612 | *lpr-1(cs249 [lpr-1::SfGFP]) I original version* | This work |
| UP3648 | *lpr-1(cs249 [lpr-1::SfGFP]) I outcrossed version* | This work |
| UP3657 | *scav-2(ok877) I; let-653(cs178) IV* | This work |
| UP3665 | *lpr-1(cs249 [lpr-1::SfGFP])) I; qxIs257[ced-1pro::NUC-1::mCherry]* | This work |
| UP3666 | *lpr-3(cs250 [SfGFP::lpr-3] X* | Cohen et al 2020 |
| UP3697 | *lpr-1(cs251 [lpr-1::mCherry]) I* | This work |
| UP3711 | *lpr-1(cs251 [lpr-1::mCherry]) I; lpr-3(cs250 [SfGFP::lpr-3] X* | This work |
| UP3745 | *lpr-1(cs251 [lpr-1::mCherry]) I; qxIs630 [scav-3::GFP]* | This work |
| UP3746 | *let-653(cs262 [LET-653::SfGFP]) IV* | Cohen et al 2020 |
| UP3808 | *lpr-3(cs266[mCherry::lpr-3]) X* | This work |
| UP3820 | *csIs87 [lpr-1pro::ss-mCherry] X; qxIs630 [scav-3::SfGFP]* | This work |
| UP3823 | *lpr-3(cs266 [mCherry::lpr-3] X; qxIs630 [scav-3::SfGFP]* | This work |

| | | |
|---|---|---|
| UP3862 | *lpr-1(cs207)/hT2[qIs48] I; lpr-3(cs250 [SfGFP::lpr-3] X* | This work |
| UP3957 | *lpr-1(cs249 [lpr-1::SfGFP]) I; lpr-3(cs266 [mCherry::lpr-3]) X* | This work |
| UP3986 | *scav-2(cs289 [scav-2::SfGFP]) III* | This work |
| UP4002 | *lpr-1(cs207)/hT2[qIs48]; scav-2 (cs289 [scav-2::SfGFP])/hT2 III* | This work |
| UP4301 | *lpr-1(cs207) I; pld-1(ok2222) II* | This work |
| UP4302 | *lpr-1(cs207) I; pld-1(ok2222) II* | This work |
| UP4354 | *lpr-1(cs207) I/ hT2[qls48] (I;III); pcyt-1(et9) X* | This work |
| UP4374 | *lpr-1(cs207)/hT2[qIs48] I; scav-2(ok877)/hT2 III; lpr-3(cs250 [SfGFP::lpr-3] X* | This work |
| UP4384 | *lpr-1(cs207)/hT2[qIs48] I; scav-2(ok877)/hT2 III; let-653(cs262 [LET-653::SfGFP]) IV* | This work |
| UP4386 | *lpr-1(cs207)/hT2[qIs48] (I;III); grl-2(syb7064 [SfGFP::GRL-2]) V* | This work |
| UP4387 | *lpr-1(cs207) I; scav-2(ok877) III; grl-2(syb7064 [SfGFP::GRL-2]) V* | This work |
| UP4391 | *lpr-1(cs207)/hT2[qIs48] (I;III); let-4(mc95 [LET-4::GFP]) X* | This work |
| UP4392 | *lpr-1(cs207)/hT2[qIs48] (I;III); let-653(cs262 [LET-653::SfGFP]) IV* | This work |
| UP4395 | *scav-2(ok877) III; lpr-3(cs250 [SfGFP::lpr-3] X* | This work |
| UP4400 | *lpr-1(cs207)/hT2[qIs48] (I;III); csIs66 [let-653pro::SfGFP::LET-653(ZP), let-653pro::PH-mCherry, lin-48pro::mRFP] X* | This work |
| UP4409 | *let-653(cs178) IV; lpr-3[cs250 [SfGFP::LPR-3]) X; csEx358[lpr-1p::LET-653b, unc-119p::GFP]* | This work |
| UP4410 | *scav-2(ok877) III; let-653(cs178) IV; lpr-3[cs250 [SfGFP::LPR-3]) X* | This work |
| XW5399 | *qxIs257 [ced-1pro::nuc-1::mCherry]* | Miao et al 2020 |
| XW8056 | *qxIs630 [scav-3::GFP]* | Miao et al 2020 |

**Table S2: Plasmids, Primers, and guide RNAs**

| Plasmids | Description |
|---|---|
| pCW11 | source of SfGFP |
| pDD162 | eft-3pro::Cas9 + U6pro:empty sgRNA |
| pJC39 | SfGFP^SEC^3xFLAG |
| pJJR83 | mCherry^SEC^3xFLAG |
| pEP75 | grl-2pro::scav-2(cDNA) |
| pEP88 | scav-2pro::NLS-GFP |
| pPD49.26 | expression plasmid backbone |
| pPD122.13 | source of NLS-GFP |
| pRFR66 | eft-3pro::Cas9 + U6pro::lpr-1sgRNA (5'-gatacggcagagtttttgac-3') |
| pRFR67 | lpr-1^SEC^SfGFP repair plasmid |
| pRFR68 | lpr-1^SEC^mCherry repair plasmid |

| Primers | Sequence |
|---|---|
| oAB4 (R) | aaaaatatatataactctctactttttttaatttgttaaaactgcaaaaattttcaaaatcaactcaaatt atcgatttttctgcaattttcaaaaattcaaggatctaattagctataTCATTTGTAGAGCTC ATCCATG |
| oAB5 (F) | AGCAAAGGAGAAGAACTTTTCAC |
| oAB6 (R) | TTTGTAGAGCTCATCCATGCC |
| oAB10 (F) | TCTGTGACCCGATTCTCGGACGGCGGTCAAAACGGGCAAATGGA TTGTCACCGGAACCTCTTCCTCCGTATGTTCCGAAGGGAGAACCA AACGGATATCACCATAATCATGCCTACATGAGCAAAGGAGAAGAA CTTTTC |
| oEP89 (F) | GACGGCTAGCAAAATGCCTTCCAGCAATAATTCG |
| oEP90 (R) | GGGAAAGAGCTCTCACATGTAGGCATGATTATGG |
| oEP92 (R) | gggaaGTCGACGAATAAAGCTTCTGTTCTCTGTCC |
| oEP96 (F) | gggaaGCATGCCCGTGCTAAATCCGAAGTATTAC |

| crRNAs | Sequence |
|---|---|
| crAB1 | ACATGTAGGCATGATTATGG |
| crAB2 | TAATTAGCTATATCACATGT |

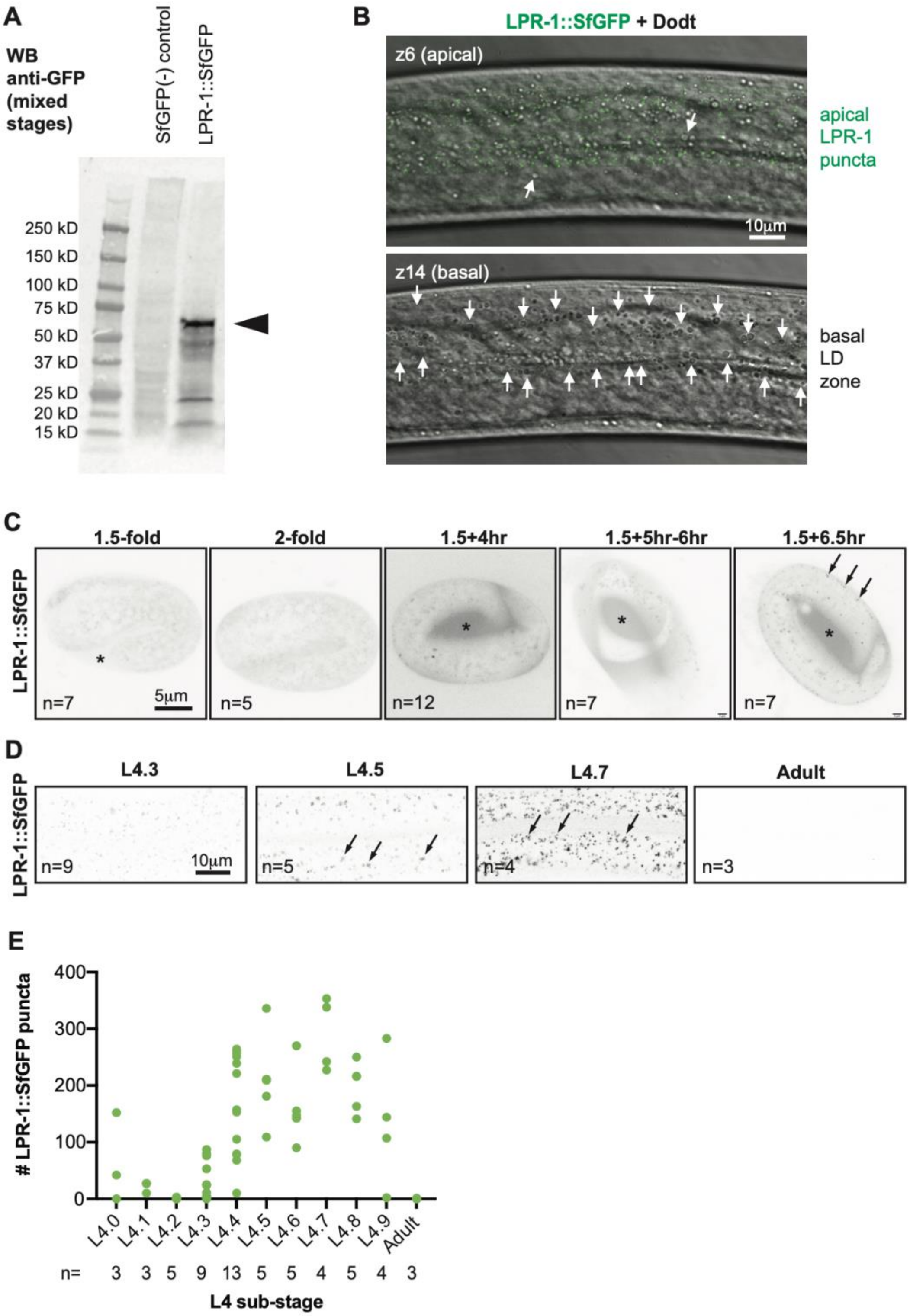

**Figure S1: LPR-1::SfGFP marks apical puncta and the extraembryonic space**

**Figure S1: LPR-1::SfGFP marks apical puncta and the extraembryonic space**
A. Western blot of mixed-stage worm lysates probed with anti-GFP antibody. The predominant band (arrowhead) matches the expected 57kD size of full-length tagged LPR-1. Image is representative of two independent blots. B. Confocal slices of the epidermis in a mid-L4 animal. LPR-1::SfGFP puncta are concentrated apically, wheres most lipid droplets (arrows) are concentrated basally within the tissue. C. Timeseries of embryonic development, showing LPR-1::SfGFP accumulating within the extraembryonic space (asterisk) and visible in some puncta (arrows). D. Timeseries of L4 larval development, showing LPR-1::SfGFP puncta (arrows). E. Quantification of puncta number across L4 stages. Puncta were counted within 61 X 38.5 micron single confocal Z-slice images of the mid-body, as shown in D.

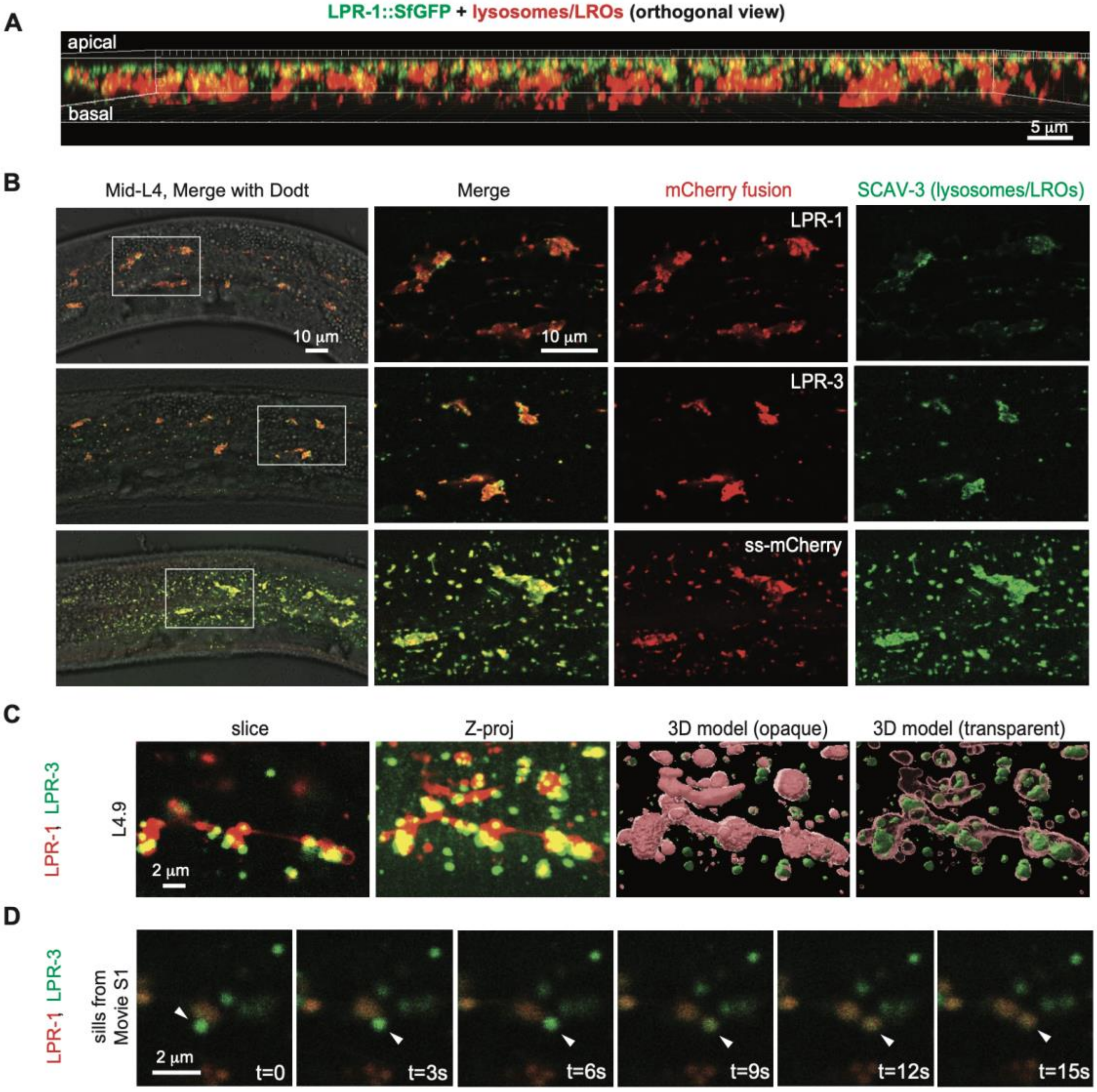

**Figure S2: LPR-1, LPR-3, and control secreted mCherry fusions accumulate in epidermal lysosomes**

A. Orthogonal (X-Z) view of confocal Z-stack, showing apical LPR-1::SfGFP puncta (green) relative to NUC-1::mCherry-marked lysosomes (red). Note that larger lysosomes are located basally in the epidermis. B. mCherry-tagged lipocalins and control fusions (red) accumulate strongly within epidermal lysosomes marked by SCAV-3::SfGFP (green). Confocal Z-slices, with boxed regions shown magnified at right. C. SfGFP::LPR-3 puncta cluster near lysosome-like structures marked by LPR-1::mCherry. 3D models were generated using Imaris software (Oxford instruments, UK). D. Stills from Movie S1, showing SfGFP::LPR-3 puncta fusing with a lysosome-like structure marked by LPR-1::mCherry. All images are representative of at least n=10 specimens imaged, except panel D which is representative of 3 movies.

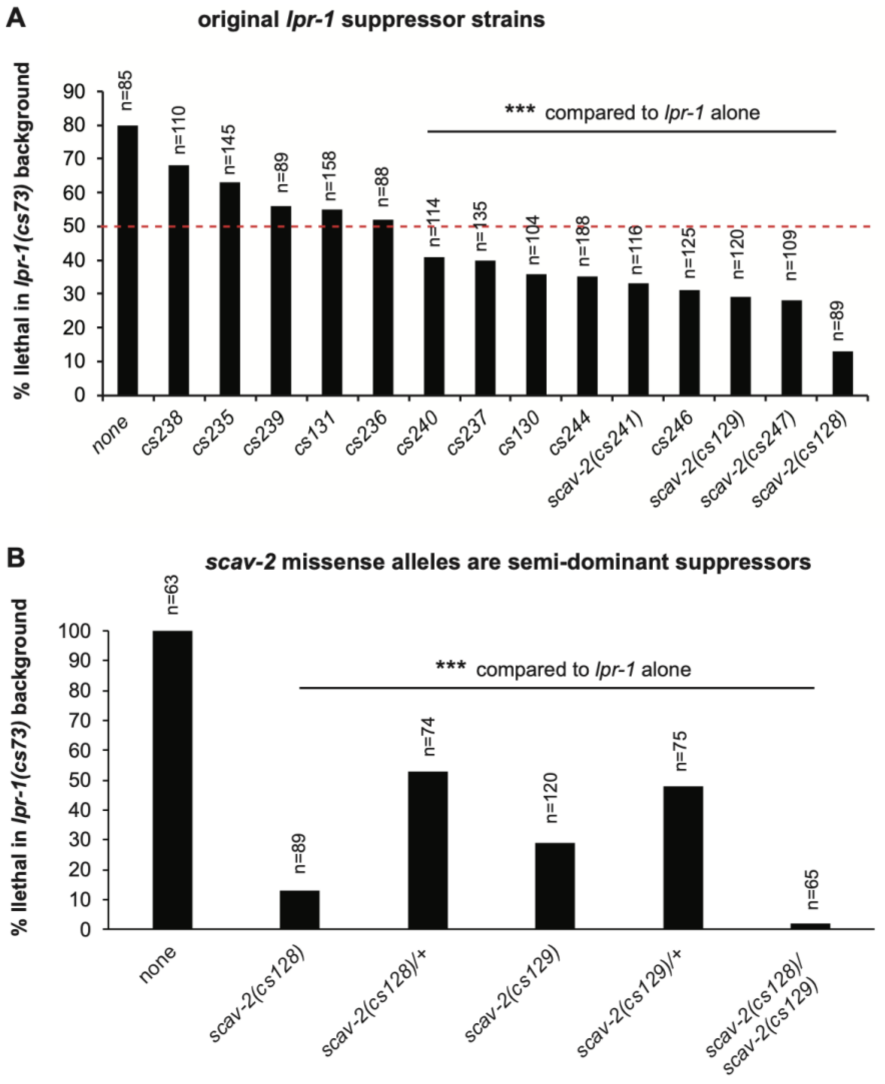

**C**

| Allele | DNA change | Protein change | First obtained as |
|---|---|---|---|
| *scav-2(cs128)* | *ccg to tcg* | P353S | *lpr-1* suppressor |
| *scav-2(cs129)* | *ccg to tcg* | P385S | *lpr-1* suppressor |
| *scav-2(cs230)* | *gga to gaa* | G349E | *let-653* suppressor |
| *scav-2(cs241)* | *gaa to aaa* | E282K | *lpr-1* suppressor |
| *scav-2(cs247)* | *aca to ata* | T343I | *lpr-1* suppressor |
| *scav-2(ok877)* | 2533 bp deletion/STOP | | deletion project allele |

**Figure S3: *lpr-1* suppressor alleles**

A. Quantification of survival in original *lpr-1* suppressor strains. ***, p<0.0001, Fisher's Exact Test, compared to parental strain. Four of the nine top suppressors contained *scav-2* alleles. B. Two *scav-2* alleles tested are each semi-dominant suppressors of *lpr-1* lethality. The data for the homozygous mutants is identical to that in A. C. Molecular lesions in *scav-2* alleles.

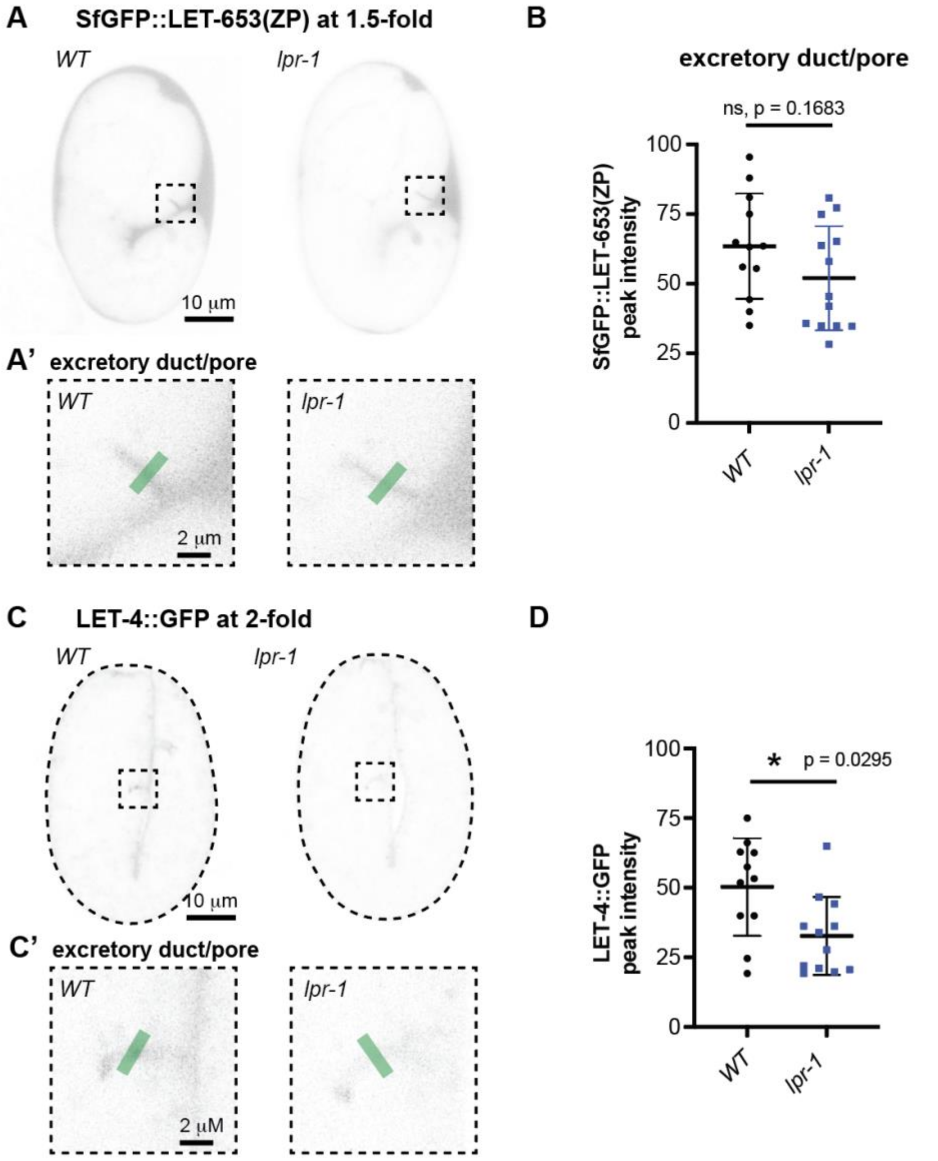

**Figure S4: *lpr-1* loss has little or no effect on excretory duct/pore matrix accumulation of LET-653 and LET-4 fusions**

A,B. Loss of *lpr-1* does not substantively impact LET-653(ZP) accumulation within the developing duct and pore tubes. A. Inverted projections of five confocal z-slices from 1.5-fold embryos. Boxed region is shown magnified in A'. Green line indicates duct/pore lumen region analyzed in B. B. Peak intensities of LET-653(ZP)::SfGFP (*csIs66*) duct/pore signal as assessed using the Plot Profile tool in FIJI. ns, Mann-Whitney U test. C-D. Loss of *lpr-1* may slightly decrease LET-4::GFP accumulation within the developing duct and pore tubes. C. Inverted projections of five confocal z-slices from 2-fold embryos. Boxed region is shown magnified in C'. Green line indicates duct/pore lumen region analyzed in D. D. Peak intensities of LET-4::GFP duct/pore signal as assessed using the Plot Profile tool in FIJI. *, p= 0.0295, Mann-Whitney U test.

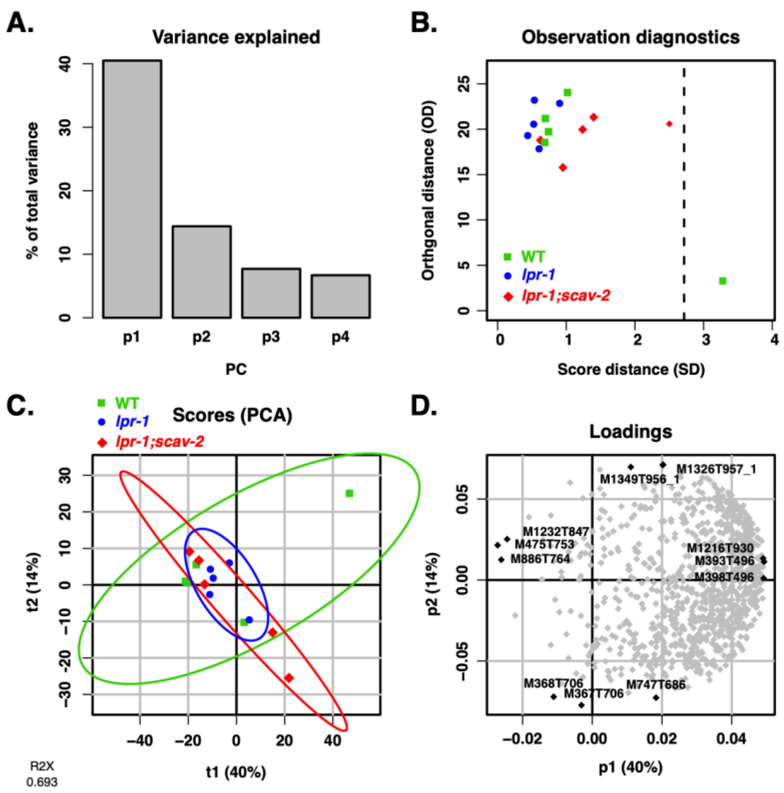

**Figure S5: *lpr-1* mutants do not have large changes in lipid abundance**

A. Principal Component Analysis (PCA) of *C. elegans* 1.5-fold embryo samples based on positive ion mode LC-MS data. The first two dimensions explain 40% and 14% of variance. B. A plot of score distance vs. orthogonal distance was used to visualize outliers. One biological sample was removed from the analysis. C. Wild-type animals (N2, green), single mutant (*lpr-1(cs207)*, blue), and double mutant (*lpr-1(cs207); scav-2(ok877)*, red) showed no consistent separation between genotypes across principal components t1 and t2, indicating no robust genotype-specific differences in overall lipid content. **D**. The most representative lipid features contributing to the principal components are visualized in the PCA loading plot.

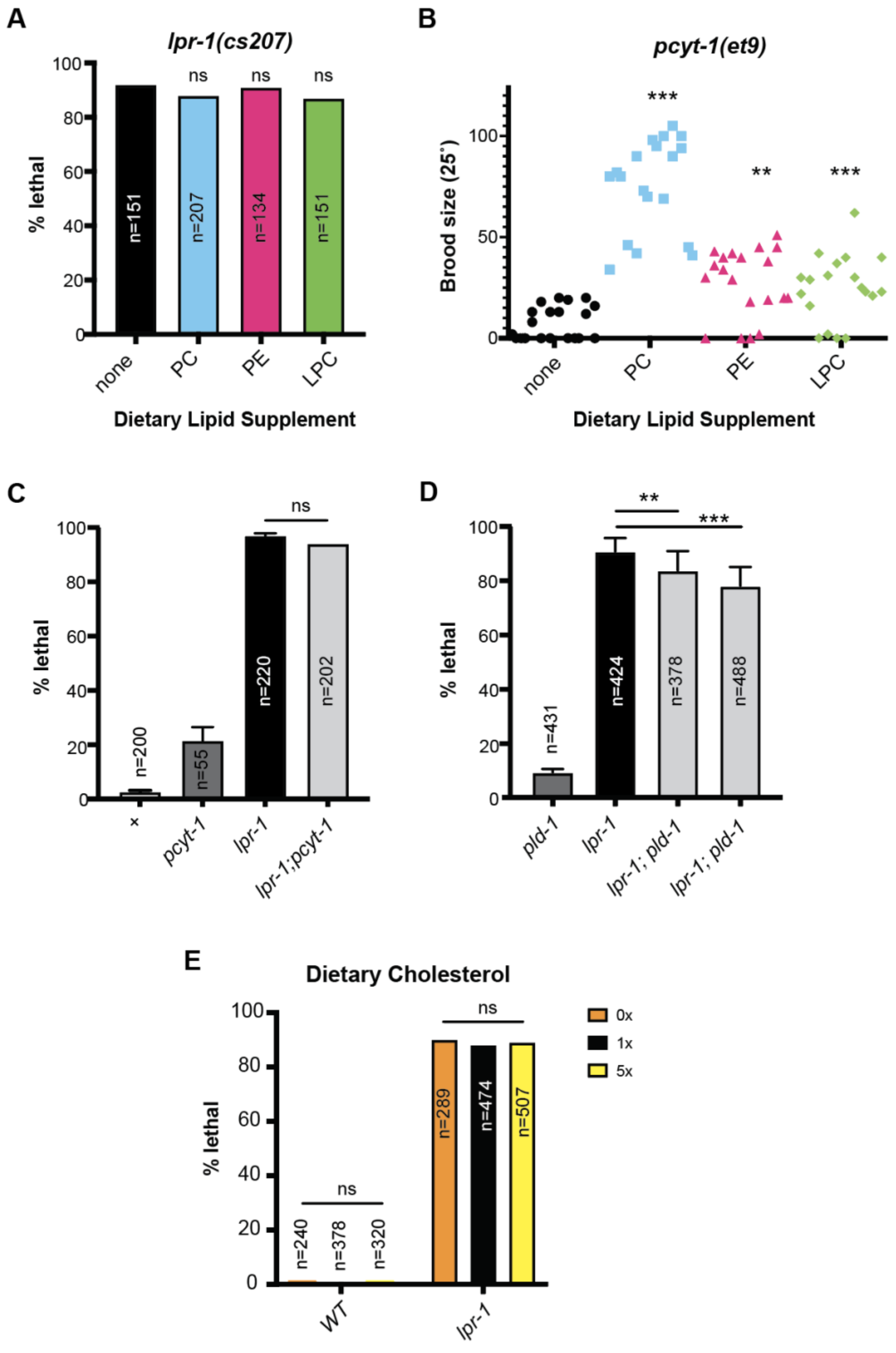

**Figure S6: Dietary phospholipid or cholesterol manipulations don't affect *lpr-1* mutant survival**

**Figure S6: Dietary phospholipid or cholesterol manipulations don't affect *lpr-1* mutant survival**

A. Dietary phospholipids phosphatidylcholine (PC), phosphatidylethanolamine (PE), and lysophoshatidylcholine (LPC) did not improve *lpr-1* survival. B. Dietary phospholipids did improve fertility of a mutant in *pcyt-1*, which encodes the *C. elegans* ortholog of phosphocholine cytidyltransferase (CCT), the rate-limiting enzyme in both the Kennedy (*de novo*) and phosphatidylethanolamine methyl transferase (PEMT) pathways for PC biosynthesis [93]. C. *pcyt-1(et9)* did not affect *lpr-1* survival. D. A null mutation in Phospholipase D, *pld-1(ok2222),* slightly improved *lpr-1* survival. The effect was seen in two independent lines generated. E. Neither increasing nor increasing dietary cholesterol affected *lpr-1* survival. Panels show pooled data from one (A,E), two (B,C) or three (D) experimental trials, with each trial containing broods of at least six worms. All data analyzed by Fisher's Exact Test relative to *lpr-1* control, except B analyzed by Mann-Whitney U Test relative to *pcyt-1* control. $^{**}p<0.001$, $^{***}p<0.0001$, ns $p>0.05$.
